# Molecular and structural characterization of lenalidomide-mediated sequestration of eIF3i

**DOI:** 10.1101/2022.08.03.502648

**Authors:** Zhi Lin, Dacheng Shen, Bo Yang, Christina M. Woo

**Affiliations:** Department of Chemistry and Chemical Biology, Harvard University, Cambridge, Massachusetts 02138 United States

**Keywords:** cereblon, lenalidomide, targeted protein degradation, chemical proteomics

## Abstract

Lenalidomide is a ligand of the E3 ligase substrate adaptor cereblon (CRBN) that achieves its clinical effects in part by promotion of substrate recruitment and degradation. In contrast to prior substrates, eIF3i is recruited but not degraded upon complex formation with lenalidomide and CRBN, although the structural details and mechanistic outcomes of this interaction are unresolved. Here, we characterize the structural basis and mechanistic outcomes of lenalidomide-induced sequestration of eIF3i from the eIF3 complex. Identification of the binding interface on eIF3i by a covalent lenalidomide probe and chemical proteomics rationalizes the sequestration event. We further connect eIF3i and CRBN to known lenalidomide effects on angiogenic markers, Akt1 phosphorylation, and associated antiangiogenesis phenotypes. Finally, we find that eIF3i sequestration is observed in MM.1S and MOLM13 cells after the degradation of other substrates, such as IKZF1. The defined binding interface elucidated by chemical proteomics, and connection of eIF3i sequestration to phenotypic outcomes of lenalidomide open future directions in designing new chemical adaptors for protein sequestration as a strategy to selectively control translational profiles and downstream cellular function.

## INTRODUCTION

Lenalidomide (Len) is an analog of thalidomide that is used to treat multiple myeloma and del(5q) myelodysplastic syndrome. Lenalidomide and its derivatives achieve their clinical activity in part by engagement of the E3 ligase substrate adaptor CRBN, which promotes the recruitment and degradation of target proteins, including IKZF1, IKZF3 and CK1α.^1-3^ The growing definition of mechanistic targets of lenalidomide has driven parallel growth in the application of CRBN ligands in targeted protein degradation strategies and the engagement of CRBN in the clinic for novel therapies.^4, 5^ Lenalidomide and its analogs additionally possess immunosuppressive, immune co-stimulatory and antiangiogenetic effects, whose mechanistic targets are only partially described.^6, 7, 8^

In our pursuit to discover additional targets of lenalidomide, we recently reported the eukaryotic translation initiation factor 3 subunit i (eIF3i) as a novel target for lenalidomide in epithelial cells using a photoaffinity labeling (PAL) and chemical proteomics strategy.^9^ eIF3i is a subunit of the eIF3 complex, which is composed of 13 subunits and involved in the protein translation initiation process, including regulation of ribosomal complexes such as 40S and 60S prior to initiation, and 43S pre-initiation complex in mRNA scanning and recognition.^10-12^ We previously found that eIF3i is stabilized in a complex with CRBN and lenalidomide, and unlike other known targets of lenalidomide, is not degraded upon recruitment to CRBN. However, the structural underpinning of the interaction, the functional outcome on the eIF3 complex, and the contribution of eIF3i recruitment to the overall effects of lenalidomide remained to be defined (**Figure 1a**). Structurally, eIF3i is composed of a helical WD40 wheel, which is unique from other recruited substrates that bear a beta-loop as the recognition motif.^13^ Loss of eIF3i may affect the functions of the eIF3 complex in translation initiation and elongation,^14^ and thus impact the regulation on angiogenic marker vascular endothelial growth factor A (VEGFA).^15^ Other reported roles of eIF3i may also be impaired, such as promoting the constitutive phosphorylation on Akt1.^16, 17^ These phenotypes associated with eIF3i loss are reminiscent of phenotypic effects that are measured for lenalidomide,^18, 19^ which may be interconnected.

**Figure 1.**
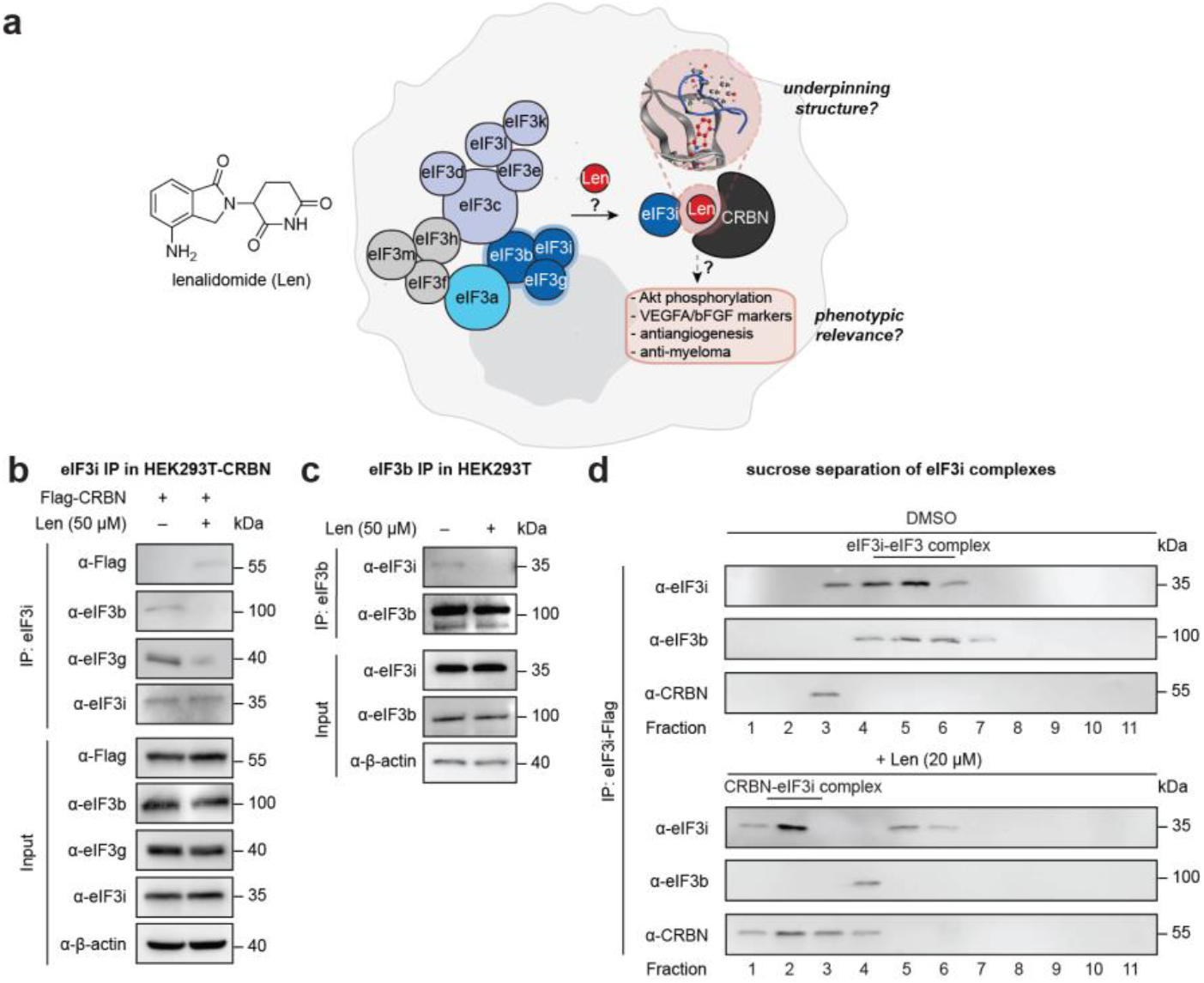
Lenalidomide sequesters eIF3i from the eIF3 complex. **(a)** Overview of the structural and mechanistic foci in this study. **(b)** Coimmunoprecipitation of endogenous eIF3i from HEK293T-CRBN cells. **(c)** Coimmunoprecipitation of endogenous eIF3i in HEK293T cells. **(d)** Representative sucrose gradient analysis of purified eIF3i-Flag complexes with or without lenalidomide treatment. Sample were separated to 11 fractions, and the eIF3i-eIF3 complex and CRBN–lenalidomide–eIF3i complex were examined by immunoblotting.

Here, we report the structural features and mechanistic outcomes of lenalidomide-mediated eIF3i recruitment. We find that eIF3i is sequestered from the eIF3 complex by lenalidomide, which is rationalized by characterization of the binding interface using an electrophilic covalent probe of lenalidomide. By using biochemical assays such as polysome profiling, ELISA assay and cellular assays, we found that sequestration of eIF3i by lenalidomide is connected to changes in the translational profiles, VEGFA sensitivity in epithelial cells, Akt1 phosphorylation, and antiangiogenetic effects in human umbilical vein endothelial cells (HUVECs). Cellular time course studies further demonstrate that the interaction of eIF3i with CRBN and lenalidomide is prevalent in different cell types including multiple myeloma MM.1S cells. Therefore, this interaction may contribute to the physiological functions of lenalidomide, especially during extended exposure time.

## RESULTS

### Lenalidomide sequesters eIF3i from the eIF3 complex

To study the effect of lenalidomide on the eIF3 complex, we initially examined the protein associations of endogenous eIF3i after coimmunoprecipitation from HEK293T cells stably expressing CRBN (HEK293T-CRBN) with or without lenalidomide at 50 µM,^9^ a concentration relevant to the anti-inflammatory and antiangiogenetic uses of lenalidomide.^18-20^ Given the direct interactions of eIF3i with eIF3b and eIF3g elucidated in the cryo-EM and crystal structure of the eIF3 subcomplex (b:g:i) in the eIF3 complex,^14, 21, 22^ we found that eIF3i coimmunoprecipitated with eIF3b and eIF3g, as expected (**Figure 1b**). However, eIF3i is reassociated with Flag-CRBN after addition of lenalidomide, and levels of eIF3b and eIF3g immunoprecipitated correspondingly decreased. Likewise, immunoprecipitation of endogenous eIF3b from HEK293T cells showed a strong interaction with eIF3i that is decreased in the presence of lenalidomide (**Figure 1c**). These data indicate that eIF3i is sequestered from the eIF3 complex upon treatment with lenalidomide.

We then quantified the disproportionation of eIF3i with the eIF3 complex or with CRBN observed on lenalidomide treatment. We performed a sucrose gradient analysis of the eIF3i-Flag immunoprecipitated from HEK293T cells to separate the complexes (**Figure 1d**). We found that eIF3i is associated with eIF3b in the untreated cells but co-elutes in two complexes in the presence of lenalidomide. After treatment with lenalidomide, the majority of eIF3i was observed in complexation with CRBN, without association to eIF3b. Protein levels of eIF3i, eIF3b, eIF3g and CRBN remain unchanged over the 48 h incubation with lenalidomide (**Figure S1**).

With endogenous CRBN engagement with lenalidomide and eIF3i discovered in HEK293T, HepG2 and HeLa cells in our previous study,^9^ we additionally found the ternary complex formation in human umbilical vein endothelial cells (HUVECs) (**Figure S2a**). Coimmunoprecipitation of endogenous eIF3i in HUVECs showed that the interaction with CRBN and lenalidomide disturbed the interaction of eIF3i with eIF3b or eIF3g (**Figure S2b**). Reciprocal immunoprecipitation of eIF3b confirmed that the eIF3b-eIF3i interaction is inhibited in the presence of lenalidomide (**Figure S2c**). Evaluation of a panel of CRBN ligands showed that sequestration of eIF3i is primarily induced by lenalidomide, and observed with pomalidomide and thalidomide to a lesser degree (**Figure S3**). By contrast, larger CRBN ligands, such as CC-885, CC-220, or dBET6 do not recruit eIF3i to a complex with CRBN (**Figure S3**). Taken together, these data demonstrate that lenalidomide selectively sequesters eIF3i from the eIF3 complex to reassociate with CRBN in epithelial cells.

### Binding interfaces of eIF3i and lenalidomide is elucidated via proteomics with a covalent probe

Intrigued by the sequestration of eIF3i with lenalidomide and CRBN, we next sought to evaluate the binding surface of the ternary complex. Previously, we reported photolenalidomide (pLen) as a probe to visualize the direct interaction of lenalidomide with CRBN and eIF3i in the ternary complex and map a binding site on CRBN; however, eIF3i was impervious to our attempts to map a binding site with photolenalidomide.^9^ Given that activity-based covalent labeling chemistries provide and alternative method for labeling proteins,^23, 24^ we designed and tested a covalent probe, termed SL1, that contains an acrylamide moiety as an electrophilic warhead for nucleophilic cysteines (**Figure 2a**).

**Figure 2.**
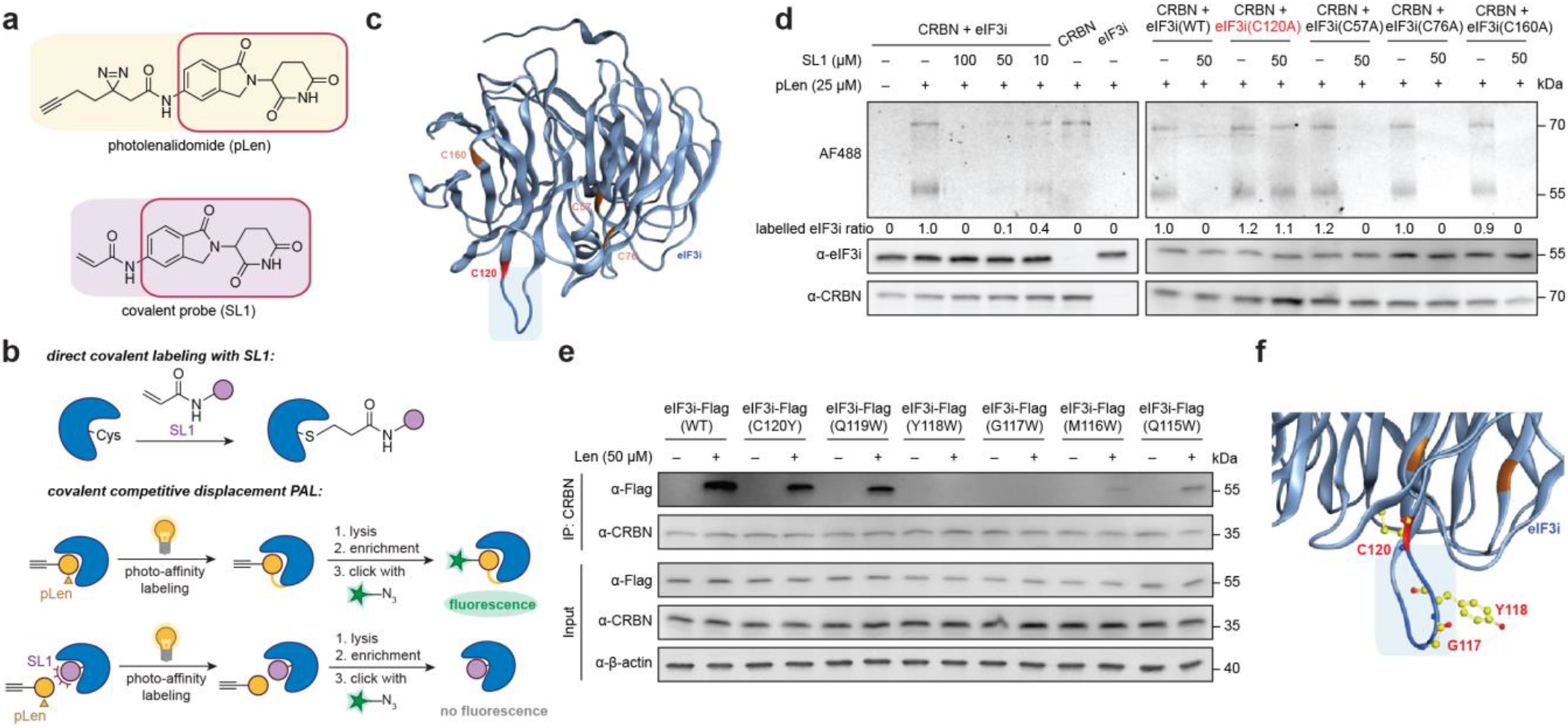
Definition of the binding interface of eIF3i and lenalidomide via chemical proteomics. **(a)** The structures of photolenalidomide and the covalent probe SL1. **(b)** Schematic demonstration of direct covalent labelling using SL1 and covalent competitive displacement photoaffinity labelling workflow using photolenalidomide and SL1. **(c)** Mutated Cys residues highlighted on the structure of eIF3i (AlphaFold: AF-Q13347-F1-model_v2). eIF3i is in light blue with the region of interest highlighted in dark blue. C120 is colored in red. Other Cys residues mutated as controls are colored in orange. **(d)** Gel-based covalent competitive displacement PAL with SL1 in competition against photolenalidomide. **(e)** Coimmunoprecipitation of wild-type (WT) eIF3i or eIF3i mutants in the presence of lenalidomide in HEK293T cells. **(f)** Proposed binding surface of lenalidomide-CRBN complex on eIF3i. eIF3i is colored in blue with residues mutated in this study (C120, Y118, G117) highlighted in yellow.

SL1 was first evaluated as a suitable covalent probe for lenalidomide (**Figure S4**). Like lenalidomide and photolenalidomide, SL1 degrades IKZF3 in MM.1S cells after 8 h in a NEDDlyation-dependent manner and promotes engagement of IKZF1 with endogenous CRBN in MM.1S cells (**Figure S4a, S4b**). When coimmunoprecipitating endogenous CRBN from HEK293T cells, SL1 enriched more eIF3i with CRBN as compared to lenalidomide and generated a stronger ternary complex in vitro with GST-eIF3i and His-CRBN by AlphaScreen (**Figure S4c, S4d**), which may indicate greater stabilization of the complex with the covalent probe and confirms the direct interaction between eIF3i and CRBN without contribution of additional cellular components.

We next focused on mapping the binding site of SL1 to eIF3i. We first performed the direct covalent labeling where recombinant His-CRBN and GST-eIF3i were incubated with 50 µM SL1 for 30 min at 37 °C before the proteins were precipitated with acetone and digested with trypsin (**Figure 2b**). By searching against His-CRBN, GST-eIF3i and common contaminant proteins, tandem MS analysis identified a cysteine residue C120 on eIF3i being labeled by SL1 (**Figure S5, Table S1**). The eIF3i residue C120 is located on the loop of the “bottom” face of the blade 3 on the WD40 wheel, which is part of the interface with eIF3b in the eIF3 complex (AlphaFold: AF-Q13347-F1-model_v2, **Figure 2c**).^25, 26^ This may explain why the interaction of eIF3i with eIF3b is impaired upon eIF3i sequestration by CRBN.

To visualize and quantify the binding event, we used photolenalidomide as a probe to visualize engagement of CRBN and eIF3i, and performed a competitive displacement of photolenalidomide using SL1 (**Figure 2b**). In brief, Flag-CRBN and eIF3i-Flag were separately purified from HEK293T cells and incubated with DMSO or different concentrations of SL1 for 30 min at 37 °C before 25 µM photolenalidomide was added and incubated for additional 30 min at 37 °C. After UV irradiation (30 s), the proteins were precipitated with acetone, resuspended, and conjugated to AlexaFluor 488 azide (AF488) for visualization by in-gel fluorescence. As expected, both eIF3i and CRBN are labeled by photolenalidomide and the labeling is competitively displaced by SL1 in a dose-dependent manner, with almost complete competition at 50 µM (**Figure 2d**). While CRBN is labeled by photolenalidomide independently, eIF3i labeling is dependent on the presence of CRBN, indicating that the interaction and labeling event is mediated by CRBN.

We then mutated eIF3i C120 and used the purified eIF3i-Flag (C120A) to evaluate whether C120 is the primary contributing residue to SL1 labeling. Three other residues were also included among the blades as controls including C57, C76 and C160 (AlphaFold: AF-Q13347-F1-model_v2, residues highlighted in orange, **Figure 2c**).^26^ The labeling ability of SL1 was tested with purified proteins similar to the aforementioned competitive displacement assay where 50 µM was chosen as the concentration for SL1 labeling (**Figure 2d**). We found that only the eIF3i C120A mutant rescued SL1 competition and is still labeled by photolenalidomide, indicating that the C120 residue is essential for SL1 engagement with CRBN and that this region of eIF3i may be critical for the binding interaction between eIF3i, lenalidomide, and CRBN (**Figure 2c**).

To model the binding interaction with eIF3i, we performed molecular docking of the bottom-face loop of the blade 3 on eIF3i where C120 resides with the lenalidomide– CRBN/DDB1 complex (PDB 4CI2) using Molecular Operating Environment (MOE, **Figure S6a**). The protein– protein docking results were scored with the GBVI/WSA dG scoring function and the ligand interactions of the docked structure with the lowest score were analyzed. We found that while the glutarimide ring of lenalidomide remains in the binding pocket of CRBN (colored in grey), the isoindolinone ring is close to the bottom-face loop of eIF3i with residue C120.

In parallel, in order to gain structural insight to the eIF3i binding surface with CRBN and lenalidomide, we performed a tryptophan-scanning mutagenesis on the bottom-face loop of blade 3 on eIF3i (residues 120-115, **Figure 2c, 2e**). We found that two eIF3i mutants, Y118W and G117W, selectively escape engagement of CRBN and lenalidomide by coimmunoprecipitation (**Figure 2e**). Mutations on M116 and Q115 also mitigated binding ability, but not mutations on C120 or Q119, possibly due to their relatively distal position from the binding pocket with lenalidomide. We further performed molecular modeling of the Y118W and G117W mutations on eIF3i (**Figure S6b**). Comparison of the overlaying structure indicates that these two point mutations introduce steric hindrance that may block binding with CRBN and lenalidomide, which is consistent with the coimmunoprecipitation results.

### eIF3i sequestration affects the functions of both eIF3i and the eIF3 complex

With a stronger structural rationalization of the sequestration of eIF3i from the eIF3 complex by lenalidomide, we next sought to explore the downstream effect of this interaction. Since eIF3i is part of the eIF3 translational complex, we first used polysome profiling to evaluate if sequestration of eIF3i will disturb the function of the eIF3 complex. Translational initiation mediated by eIF3 starts by promoting the joining to 40S with 60S to form the 80S ribosome.^10^ Previous studies on the eIF3 complex also concluded that eIF3 is involved in the elongation process, which contributes to polysome generation.^27^ Using a sucrose density gradient with HEK293T whole cell lysates with and without lenalidomide treatment, we found an increase in the 80S monosome peak and a slight decrease in the 40S and 60S complexes when overlaying the lenalidomide-treated profile with the control profile (**Figure 3a**). Polysome peaks, which are indicative of well-translated mRNAs, were slightly decreased, suggesting a change in the translation initiation:elongation ratio driven by lenalidomide treatment. mRNA levels of eIF3i and eIF3b was confirmed unchanged both in HEK293T cells and in HUVECs (**Figure S7**).

**Figure 3.**
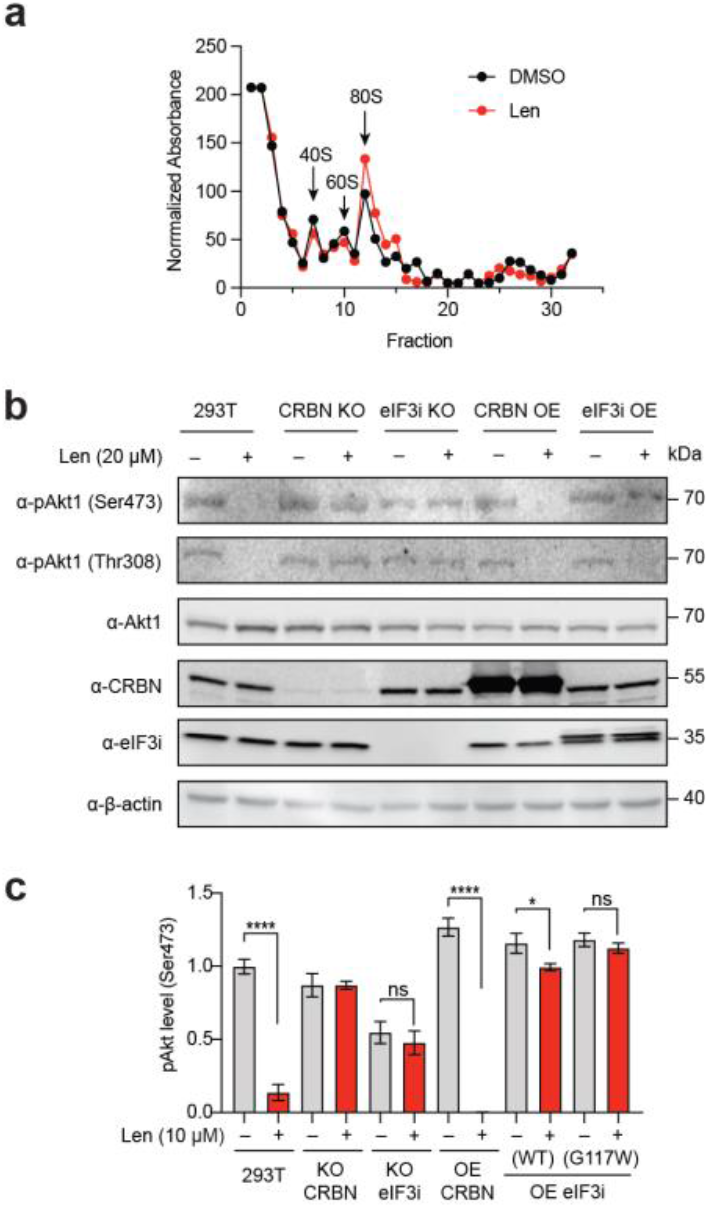
eIF3i sequestration changes the functions of eIF3i and the eIF3 complex. **(a)** Polysome profiling of HEK293T cells with or without lenalidomide analyzed by sucrose gradient analysis. The migration of the 40S, 60S and 80S ribosomes is indicated by arrows. The elution profiles of the indicated fractions were determined by absorbance at 280 nm and normalized. **(b)** Akt1 phosphorylation levels (Ser473, Thr308) with or without lenalidomide treatment by immunoblotting. **(c)** Akt1 phosphorylation level (Ser473) with or without lenalidomide treatment by ELISA assay.

In addition to participating in translation as a part of the eIF3 complex, eIF3i possesses other reported functions, such as promoting constitutive Akt1 activation.^16^ We evaluated the phosphorylation of Akt1 on both Ser473 and Thr308 by immunoblotting. Lenalidomide inhibited Akt1 phosphorylation as expected.^18, 19^ We found that lenalidomide inhibited phosphorylation of Akt1 in HEK293T cells in a manner that is dependent on CRBN and eIF3i (**Figure 3b**). We further quantified Akt1 phosphorylation on Akt1 Ser473 by ELISA and found that the inhibition of pAkt1 is dose-dependent in HEK293T cells over 1–10 µM concentrations of lenalidomide, which was similarly dependent on CRBN and eIF3i levels (**Figure 3c, Figure S8a**). As expected, CRBN overexpression (OE) enhanced the response and eIF3i OE rescued Akt1 phosphorylation. eIF3i mutant identified in the structural studies (G117W) does not engage lenalidomide in the ternary complex, thus not responding to lenalidomide treatment. These results indicate that the decrease in Akt1 phosphorylation on lenalidomide treatment is mediated by both CRBN and eIF3i. Validation of knock-out (KO) and OE for CRBN and eIF3i was demonstrated by qRT-PCR (**Figure S8b**).

### eIF3i sequestration is associated with lenalidomide antiangiogenesis in cells

We next evaluated the phenotypic relevance of the eIF3i sequestration to the CRBN–lenalidomide–eIF3i complex. Since selective ternary complex formation was observed in HUVECs, which is a cellular model for angiogenesis, we next evaluated if eIF3i and Len-mediated eIF3i sequestration is relevant to lenalidomide antiangiogenesis.

To evaluate whether the antiangiogenic properties of lenalidomide may be due to eIF3i sequestration, we measured a proangiogenic factor, the vascular endothelial growth factor A (VEGFA) in HEK293T cells (**Figure 4a**). Previous studies have established that upon lenalidomide treatment, VEGFA levels decrease.^18, 28^ We found that lenalidomide significantly decreased VEGFA mRNA levels in HEK293T cells, in a manner that was CRBN- and eIF3i-dependent (**Figure 4a**). A similar dependence on CRBN and eIF3i was observed with expression of another angiogenesis marker, the basic fibroblast growth factor (bFGF), which is also inhibited in response to lenalidomide treatment (**Figure S9**).

**Figure 4.**
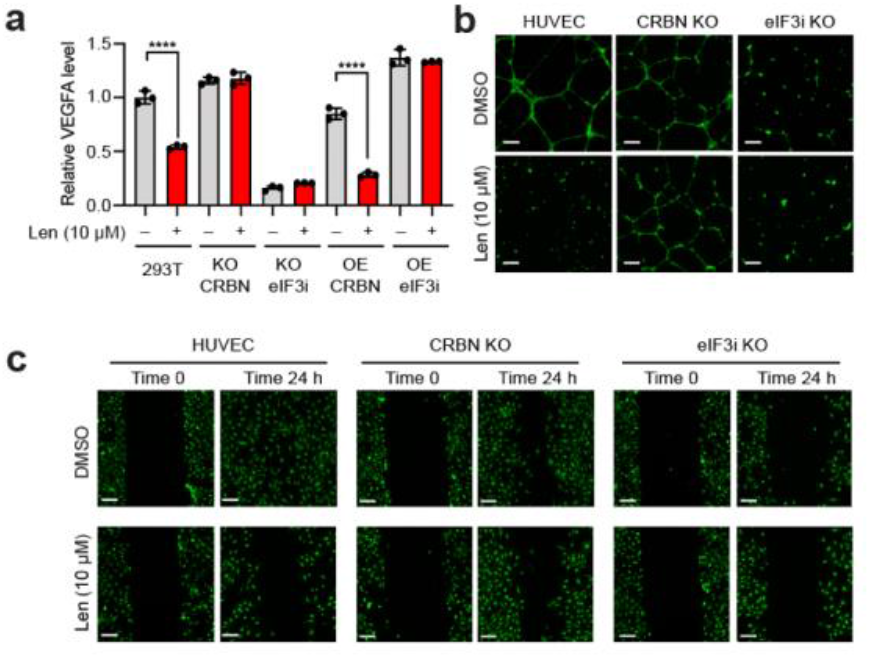
Lenalidomide and eIF3i are associated with antiangiogenesis effects. **(a)** VEGFA mRNA levels in HEK293T cells with or without lenalidomide by qRT-PCR. **(b)** Fluorescent images of tube formation assay with HUVECs with or without lenalidomide treatment for 24 h. **(c)** Fluorescent images of wound healing migration assay with HUVECs with or without lenalidomide for 24 h.

We then evaluated whether the antiangiogenetic phenotypes of lenalidomide are also dependent on both CRBN and eIF3i and performed both two-dimensional (2D) cell migration and three-dimensional (3D) cell invasion assays with HUVECs to measure angiogenesis. As a vital process for cancer cell proliferation, angiogenesis involves multiple steps including endothelial cell migration, invasion, and differentiation into capillaries.^29^ HUVEC differentiation was first confirmed with capillary tube formation of HUVECs in a 3D Matrigel matrix culture environment (**Figure 4b, Figure S10c**). We found that the tube formation ability of HUVECs was significantly decreased in the presence of lenalidomide (**Figure 4b**). HUVECs were not sensitive to lenalidomide treatment when CRBN was knocked-out (KO), while eIF3i KO resembled lenalidomide-treated WT-HUVECs. In order to visualize HUVEC invasiveness, we performed a 3D transwell assay where HUVECs were allowed to invade through Boyden chambers (**Figure S10a, Figure S10c**). The number of invasive cells that passed through the transwell membrane was quantified and compared (**Figure S10b**). Inhibition of HUVECs invasiveness by lenalidomide was similarly dependent on eIF3i and CRBN. Furthermore, we performed a wound healing assay where a monolayer of HUVECs was scratched and allowed to migrate over a period of 24 h (**Figure 4c, Figure S10c**). CRBN KO decreased the responsiveness to lenalidomide inhibition significantly, indicating that CRBN is necessary for the antiangiogenesis properties of lenalidomide. With eIF3i was KO, the migration ability of HUVECs was also inhibited. Taken together, these data suggest that lenalidomide exerts antiangiogenetic functions in HUVECs in a CRBN- and eIF3i-dependent manner.

### eIF3i is a direct target of lenalidomide in MM.1S cells after degradation of other targets

Finally, we returned to evaluate the sequestration of eIF3i in hematopoietic cell lines. Previously, we observed the sequestration of eIF3i in epithelial cell lines, but not in multiple myeloma MM.1S cells, potentially due to the stronger recruitment of IKZF1 to CRBN by lenalidomide.^9^ However, as cytotoxicity starts at 5 d incubation of lenalidomide while IKZF1 is long depleted by the CRBN– lenalidomide complex,^2, 9, 30^ we hypothesized that the proteome without lenalidomide target IKZF1 can serve as a new proteome, where subsequent complexation with eIF3i may occur.

We first monitored the protein levels of IKZF1, IKZF3 and eIF3i in MM.1S cells over time (**Figure S11**). Levels of IKZF1 and IKZF3 significantly decreased starting at 4 h and are almost completely depleted after 8–12 h, while eIF3i levels were steady up to 48 h. We therefore compared the interactions of CRBN by coimmunoprecipitation in the presence of lenalidomide at 0 h and 24 h of lenalidomide incubation in MM.1S cells, which represent time points before and after IKZF1 degradation (**Figure 5a, Table S1**). As expected, IKZF1 was significantly coimmunoprecipitated in the presence of lenalidomide [enrichment ratio (Len/DMSO) = 2.746] in MM.1S cells prior to the degradation of IKZF1. By contrast, enrichment of IKZF1 was decreased after 24 h [enrichment ratio (Len/DMSO) = 1.258], and instead eIF3i was one of the most enriched proteins [enrichment ratio (Len/DMSO) = 7.411]. We also observed histone H2A enrichment in both datasets. The preferential immunoprecipitation of IKZF1 at 0 h and eIF3i after 24 h incubation was further validated by Western blot (**Figure S12**). These data indicate that CRBN favors a ternary complex with IKZF1 in the presence of lenalidomide, but as protein levels shift in response to degradation, complexation with eIF3i can subsequently accumulate.

**Figure 5.**
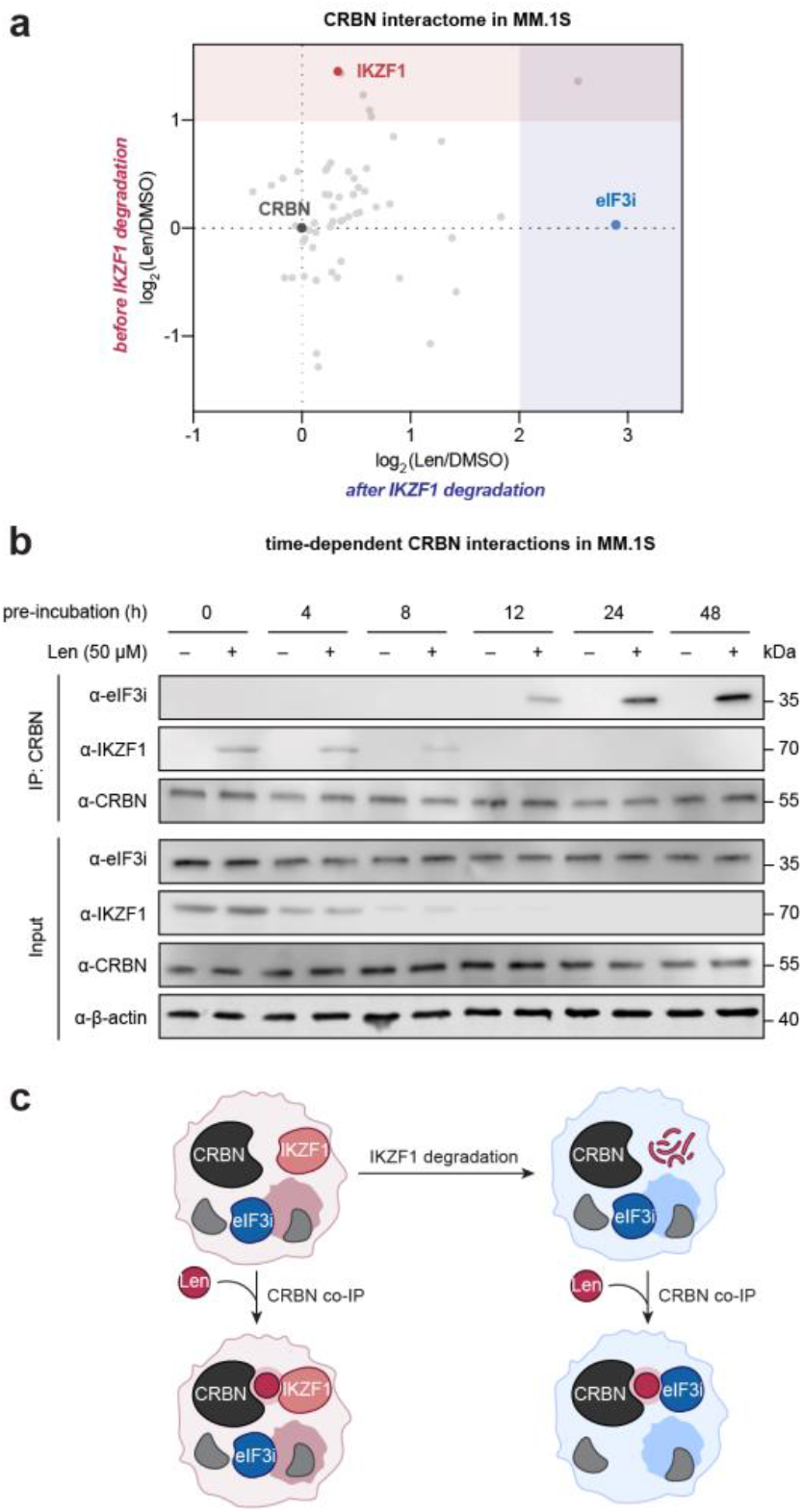
eIF3i complexes with CRBN on lenalidomide treatment in MM.1S cells after the degradation of other substrates. **(a)** CRBN interactome before IKZF1 degradation (time = 0 h) or after IKZF1 degradation (time = 48 h, 10 µM lenalidomide) by coimmunoprecipitation of endogenous CRBN with 50 µM lenalidomide. The significantly enriched region of proteins from MM.1S at time = 0 h is shaded in red. The significantly enriched region of proteins from MM.1S at time = 24 h is shaded in blue. **(b)** Endogenous CRBN coimmunoprecipitation with or without lenalidomide in MM.1S before and after lenalidomide-mediated target degradation. **(c)** Schematic model of CRBN–lenalidomide– eIF3i interaction in MM.1S when IKZF1 is degraded.

In order to validate this finding, we further performed a time-course coimmunoprecipitation experiment where the MM.1S cells were treated with lenalidomide and endogenous CRBN was immunoprecipitated from each sample (**Figure 5b**). As expected, IKZF1 levels decreased over the first 8 h while eIF3i and CRBN levels were largely unchanged over the 48 h experiment. Similarly, IKZF1 coimmunoprecipitated with CRBN in the presence of lenalidomide over the first 8 h, albeit in proportion to the decreasing levels of IKZF1. By contrast, eIF3i coimmunoprecipitated with CRBN to a greater degree over time. This shift in complexation is observable after 12 h, and appears to plateau between 24–48 h. This shift in complexation as a function of time was also observed in MOLM13 cells, which also express IKZF1 (**Figure S13**). Coimmunoprecipitation of CRBN recruited IKZF1 in a lenalidomide-dependent manner, while following depletion of these proteins using lenalidomide at 24 h, eIF3i is instead recruited to the complex. Taken together, these data indicate that eIF3i is a direct target of CRBN and lenalidomide in MM.1S and MOLM13 cells, and that this interaction becomes observable when other targets such as IKZF1 are degraded (**Figure 5c**). Although the degradation of IKZF1 is almost complete at 8 h in both MM.1S and MOLM13 (**Figure S11, S13**), 5-day incubation with lenalidomide was reported to be needed for antiproliferative effects in MM.1S and MOLM13,^2, 9, 30^ which indicates that additional factors may contribute to the cytotoxicity of lenalidomide.

## DISCUSSION

Here we report the functional, structural, and phenotypic outcomes of eIF3i sequestration by lenalidomide to a complex with CRBN. We find that the recruitment of eIF3i by CRBN and lenalidomide dissociates eIF3i from the eIF3 complex, which affects translational initiation and elongation. We developed a new covalent probe SL1, to reveal the binding surface on eIF3i that interacts with lenalidomide and CRBN in a ternary complex. We further elucidated that sequestration of eIF3i is associated with the antiangiogenic effects of lenalidomide treatment in epithelial cells, such as HEK293T cells and HUVECs, and may play a role in longer dosing regimens for MM.1S or MOLM13 cells, where recruitment of eIF3i to CRBN by lenalidomide increases over longer treatment times due to degradation of other targets like IKZF1. These findings highlight the prevalence of eIF3i engagement with CRBN in the presence of lenalidomide in different cell types. Examination of the recruitment and stabilization of substrates by other lenalidomide analogs or bifunctional degraders after the degradation of desired targets may reveal additional targets and mechanisms.

The successful identification of the binding interface between eIF3i, lenalidomide, and CRBN with the covalent probe SL1 indicates that distinct structural features exist beyond the beta-loop commonly recognized in other chemically-induced substrates.^13^ The chemical proteomics approached adapted in this study can also be useful to map the binding interface of CRBN binding targets with undefined structures, including ARID2 and p63.^31, 32^ Since the interaction with eIF3i is generalizable across different cell lines, future efforts may identify other targets of lenalidomide or derivatives that share homology with the WD40 wheel structure found in eIF3i. Chemical proteomics may enable the discovery these targets, where PAL-based chemical proteomics and activity-based chemical proteomics can be complementary. Additional studies to reveal why eIF3i is not degraded upon engagement or to engineer an eIF3i that is degradable may also open new directions in precise control of eukaryotic translation. Development of novel eIF3i-selective chemical moieties can also serve as an alternative strategy to alter transcriptional regulation. Given our observation that lenalidomide can induce interaction with eIF3i in MM.1S cells, eIF3i-selective compounds would enable further study on whether the interaction with eIF3i may be contributing to phenotypes in multiple myeloma and what direct translational outcome is associated to eIF3i sequestration.

## ASSOCIATED CONTENT

### Supporting Information

The Supporting Information is available free of charge on the ACS Publications website.

Supplementary figures, uncropped data, synthetic schemes and materials and methods (PDF)

Complete list of proteins and peptide spectral matches from proteomics experiments (XLSX)

## AUTHOR INFORMATION

Corresponding Author

* ORCID

Christina M. Woo: 0000-0001-8687-9105

Funding Sources

Support from the Ono Pharma Foundation (C.M.W.), Sloan Research Foundation (C.M.W.), Camille–Dreyfus Foundation (C.M.W.), Blavatnik Accelerator at Harvard University (C.M.W.), and Harvard University is gratefully acknowledged.

## Notes

The authors declare no competing financial interest.

## ACKNOWLEDGMENT

We thank Y. Amako, Y. Ge, W. Xu for insightful discussions; B. Budnik, R. A. Robinson, and J. X. Wang Harvard University Mass Spectrometry and Proteomics Resource Laboratory; J. Yu from Mazeed lab; C. M. Whilden from Whipple lab; T. J. Chang and C. B. Hartmann from Bauer Core Facility; E. Cronin-Furman with Olympus for technical support.

## Table of Contents

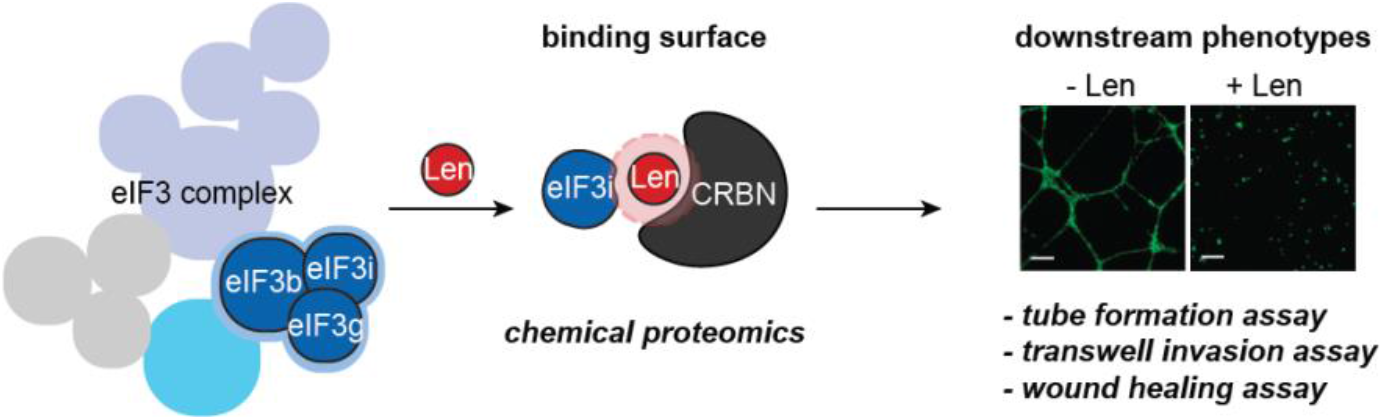

