## Supplementary material for "Molecular and structural characterization of lenalidomide-mediated sequestration of eIF3i": SI

### Index


Additional Experimental Information ........................................................................ S3

**Figure S1** Protein levels of eIF3i, eIF3b, eIF3g in HEK293T cells over lenalidomide incubation.
**Figure S2** Lenalidomide sequesters eIF3i from the eIF3 complex to form a ternary complex with CRBN in HUVECs.
**Figure S3** Coimmunoprecipitation of endogenous CRBN in the presence of lenalidomide or other common molecular glues in HEK293T cells.
**Figure S4** Validation of SL1 as a probe for lenalidomide functions
**Figure S5** Representative assignment of a conjugated eIF3i peptide with SL1.
**Figure S6** Validation of binding surface of CRBN–lenalidomide–eIF3i complex.
**Figure S7** eIF3i and eIF3b mRNA levels do not change upon lenalidomide treatment.
**Figure S8** Lenalidomide-mediated Akt1 phosphorylation decrease is associated with eIF3i and CRBN functions.
**Figure S9** Angiogenic marker bFGF levels in HEK293T cells with or without lenalidomide.
**Figure S10** Lenalidomide antiangioegensis is associated with eIF3i and CRBN functions.
**Figure S11** Validation of CRBN interactome in **Figure 1b** by immunoblotting.
**Figure S12** Time-course degradation assay of all targets in MM.1S cells.
**Figure S13** eIF3i is a direct target of lenalidomide in MOLM13 cells upon degradation of other targets.
**Figure S14 |** Uncropped immunoblots related to **Figure 1.**
**Figure S15 |** Uncropped immunoblots related to **Figure 2.**
**Figure S16 |** Uncropped immunoblots related to **Figure 3** and **Figure 5.**

**Scheme S1 |** Synthetic scheme for SL1.
**Table S1 |** Unique SL1-modified peptide assignments identified by LC-MS/MS.
**Table S2 |** Complete protein list identified by LC-MS/MS in **Figure 5a**.

General methods and instrumentation ................................................................. S20
General chemical methods and instrumentation................................................. S20
Synthetic procedures ........................................................................................ S22
Antibodies and biological reagents.................................................................... S23
Cell culture and transfection ............................................................................. S23
Plasmids and subcloning ................................................................................... S23
Immunoprecipitation assays in live cells ........................................................... S24
Sucrose gradient centrifuge fractionation assay ................................................ S25
AlphaScreen assay for binding interaction analysis........................................... S25
Direct covalent labeling with SL1 and MS sample preparation.......................... S25
Mass spectrometry sample processing for covalent binding site identification .................. S25
Mass spectrometry data analysis procedure for covalent binding site identification........... S26
Covalent competitive displacement PAL assay ................................................. S26
Structural molecular modeling ........................................................................... S26
Polysome profiling............................................................................................. S26

RNA isolation and qRT-PCR experiment ..... S27
Akt1 phosphorylation ELISA assay ..... S27
Wound healing assay ..... S27
Transwell migration assay with Boyden chambers ..... S27
Tube formation assay ..... S27
Quantitative mass spectrometry sample preparation ..... S27
High pH separation and quantitative mass spectrometry analysis ..... S28
Quantitative mass spectrometry data analysis ..... S28
Software ..... S29
Statistical analysis ..... S29
References ..... S30
NMR and IR spectrum of compound SL1 ..... S31

### SUPPORTING INFORMATION FIGURES

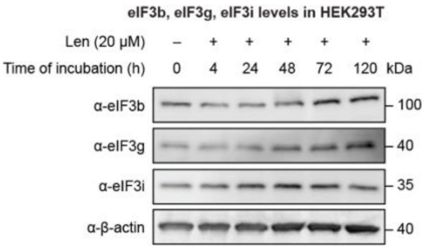


**Figure S1** Protein levels of eIF3i, eIF3b, eIF3g in HEK293T cells over lenalidomide incubation.

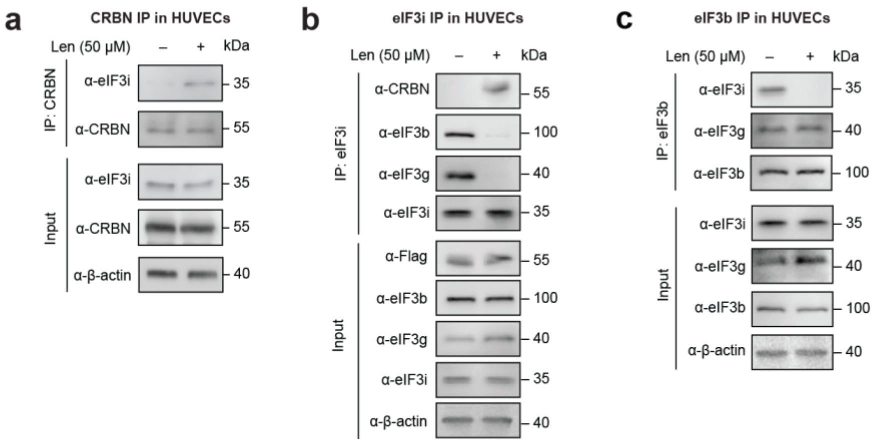


**Figure S2** Lenalidomide sequesters eIF3i from the eIF3 complex to form a ternary complex with CRBN in HUVECs. **(a)** Coimmunoprecipitation of endogenous CRBN with or without lenalidomide in HUVECs. **(b)** Coimmunoprecipitation of endogenous eIF3i with or without lenalidomide in HUVECs. **(c)** Coimmunoprecipitation of endogenous eIF3b with or without lenalidomide in HUVECs.

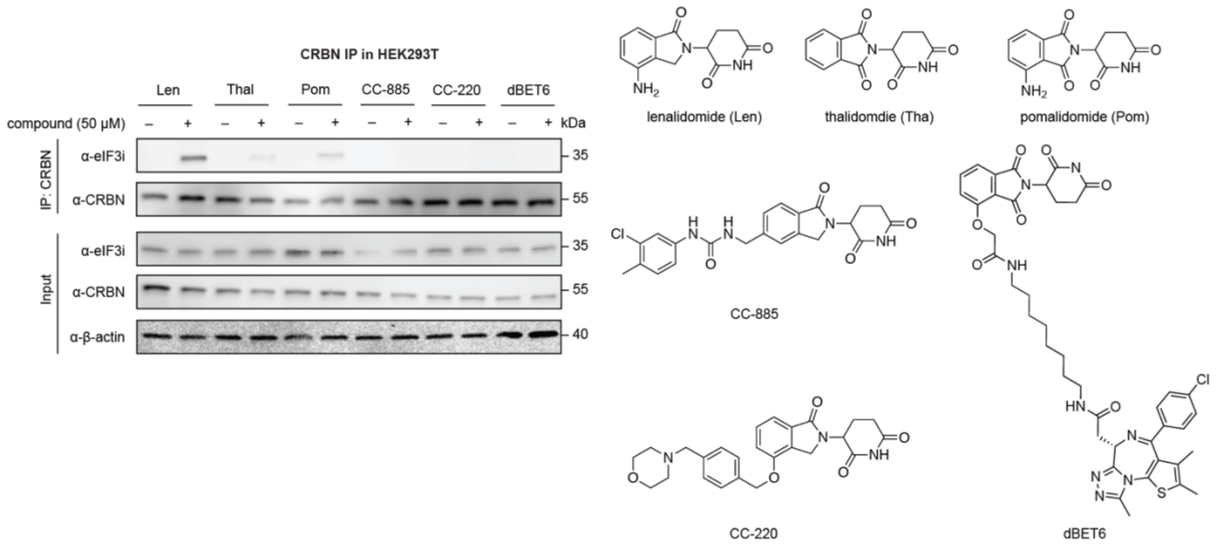


**Figure S3** Coimmunoprecipitation of endogenous CRBN in the presence of lenalidomide or other common molecular glues in HEK293T cells.

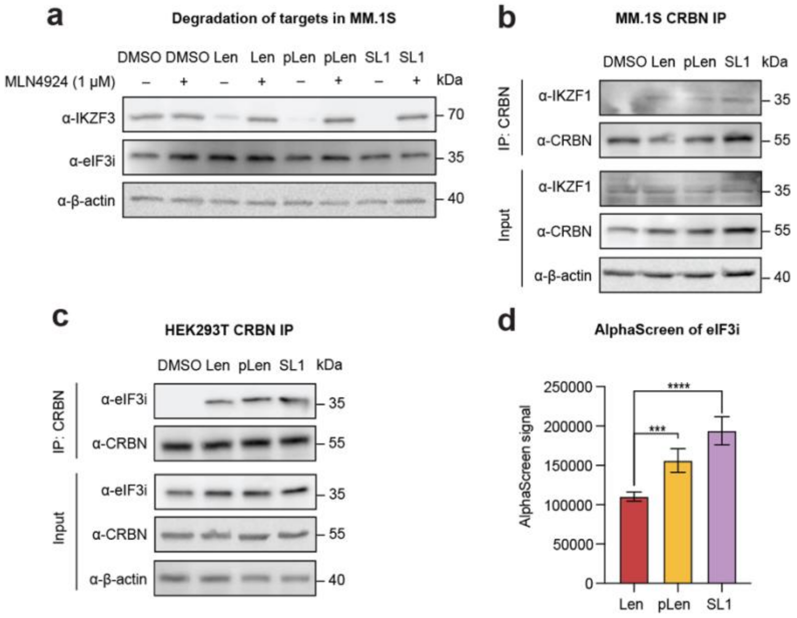


**Figure S4** Validation of SL1 as a probe for lenalidomide functions. **(a)** Western blots of substrates IKZF1 and IKZF3 in MM.1S cells after treatment with lenalidomide, photolenalidomide and SL1 for 8 h. **(b)** Coimmunoprecipitation of endogenous CRBN in MM.1S cells with lenalidomide, photolenalidomide and SL1. **(c)** Coimmunoprecipitation of endogenous CRBN in HEK293T cells with lenalidomide, photolenalidomide and SL1. **(d)** In vitro binding assay of CRBN and eIF3i induced by lenalidomide, photolenalidomide or SL1 via AlphaScreen assay.

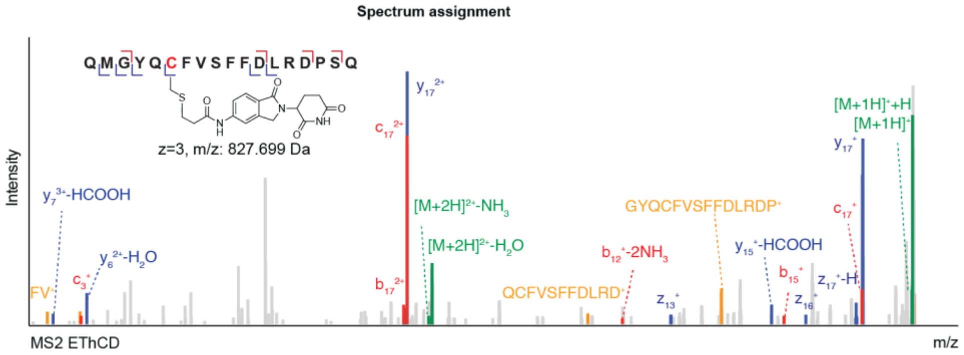


**Figure S5** Representative assignment of a peptide on eIF3i conjugated with SL1.

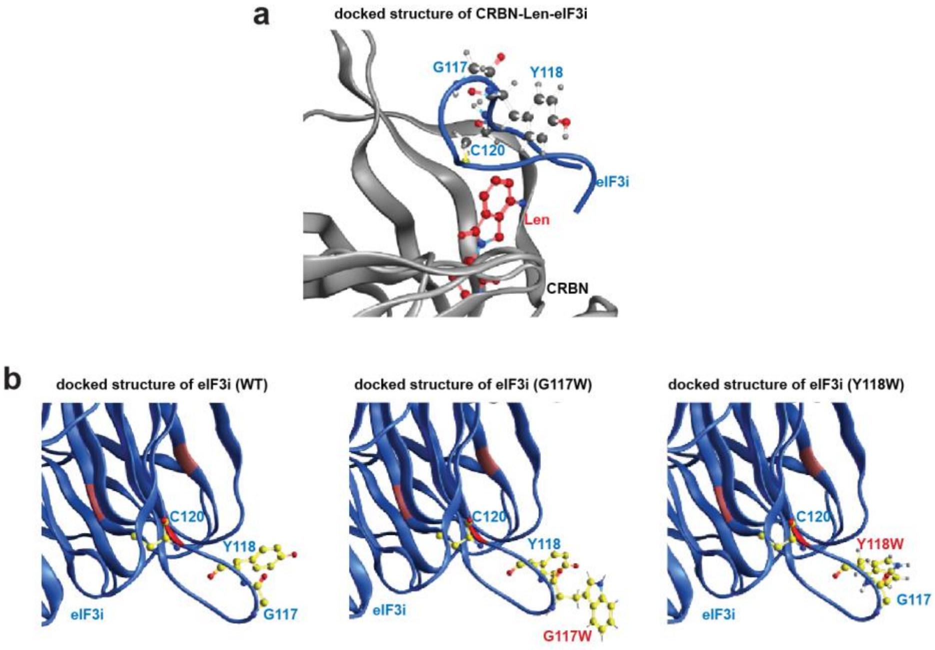


**Figure S6** Validation of binding surface of CRBN–lenalidomide–eIF3i complex. **(a)** Docked structure of the CRBN–lenalidomide–eIF3i complex. CRBN is colored in grey. eIF3i is colored in blue with residues mutated in this study highlighted. Lenalidomide is colored in red. **(b)** Docked structures of eIF3i (WT), eIF3i mutant (G117W) and eIF3i mutant (Y118W). eIF3i is colored in blue with site C120, Y118 and G117 highlighted.

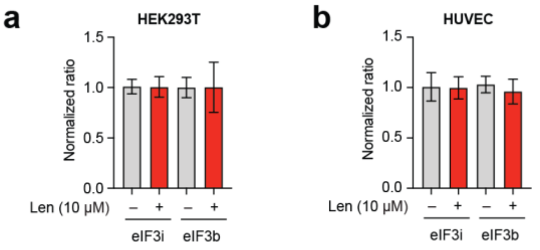


**Figure S7** eIF3i and eIF3b mRNA levels do not change upon lenalidomide treatment. **(b)** Quantification of eIF3i and eIF3b mRNA levels in (a) HEK293T cells or (b) HUVECs with or without lenalidomide by qRT-PCR.

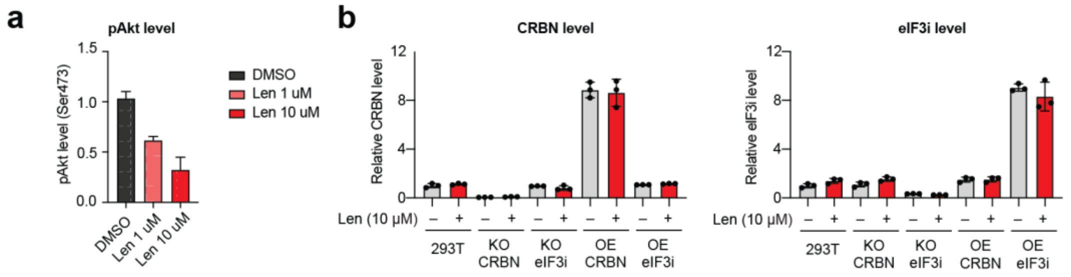


**Figure S8** Lenalidomide-mediated Akt1 phosphorylation decrease is associated with eIF3i and CRBN functions. **(a)** Dose-dependent Akt1 phosphorylation (Ser473) in HEK293T cells with different concentrations of lenalidomide. **(b)** Validation of CRBN and eIF3i knock-out (KO) and over-expression (OE) in HEK293T cells by qRT-PCR.

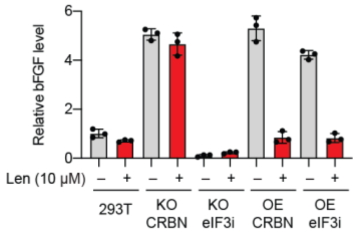


**Figure S9** Angiogenic marker bFGF levels in HEK293T cells with or without lenalidomide.

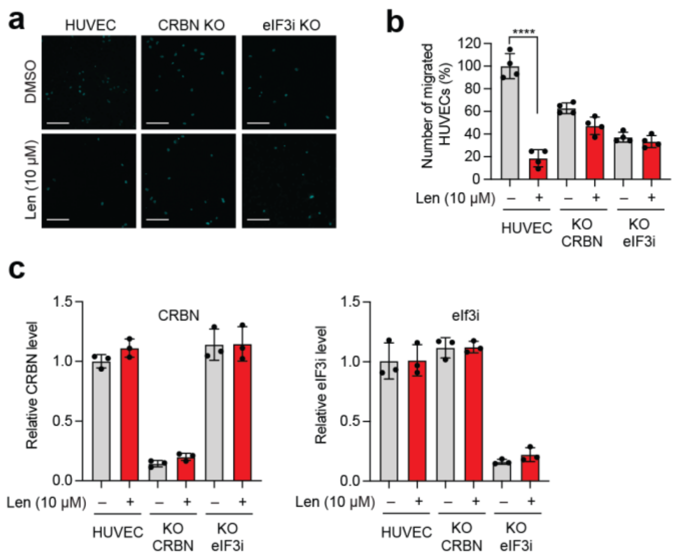


**Figure S10** Lenalidomide antiangioegensis is associated with eIF3i and CRBN functions. **(a)** Fluorescent images of migrated HUVECs with or without lenalidomide for 24 h in transwell migration assay. **(b)** Phenotypic quantification of HUVEC cell migration assay with or without lenalidomide for 24 h. **(c)** Validation of CRBN and eIF3i knock-out (KO) in HUVECs by qRT-PCR.

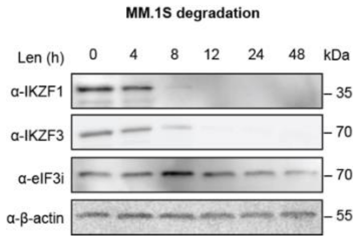


**Figure S11** Time-course degradation assay of all targets in MM.1S cells.

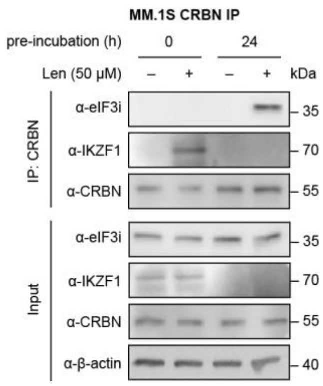


**Figure S12** Validation of CRBN interactome in **Figure 5a** by immunoblotting.

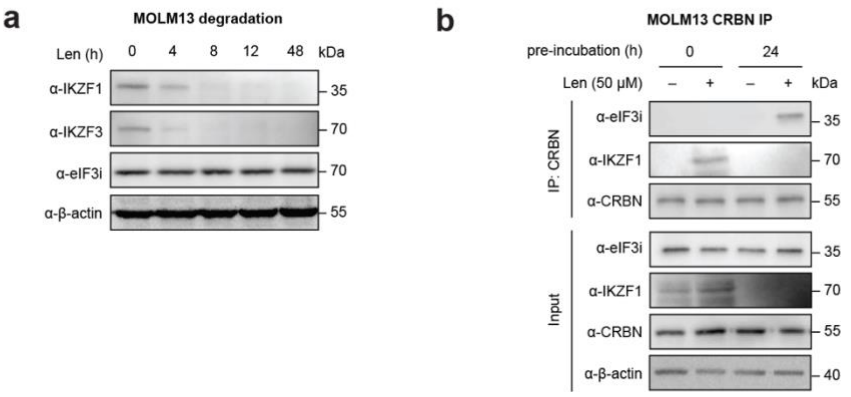


**Figure S13** eIF3i is a direct target of lenalidomide in MOLM13 cells upon degradation of other targets. **(a)** Time-course degradation assay of all targets in MOLM13 cells. **(b)** Coimmunoprecipitation of endogenous CRBN with or without lenalidomide in MOLM13 cells before and after lenalidomide-mediated target degradation.

**Figure 1b**

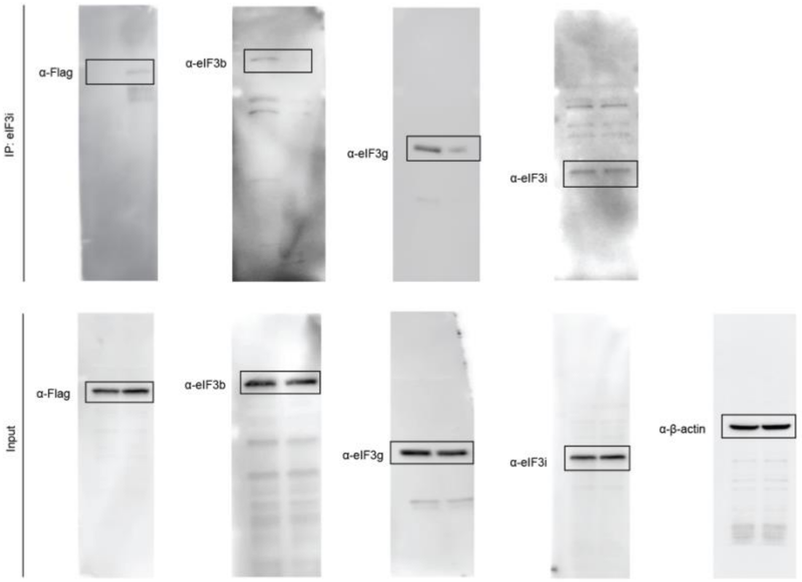


**Figure 1c**

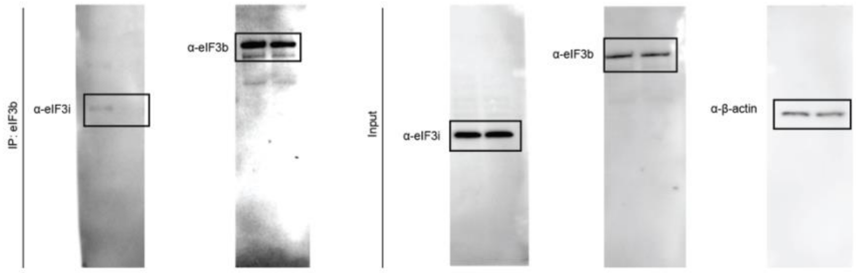


**Figure 1d**

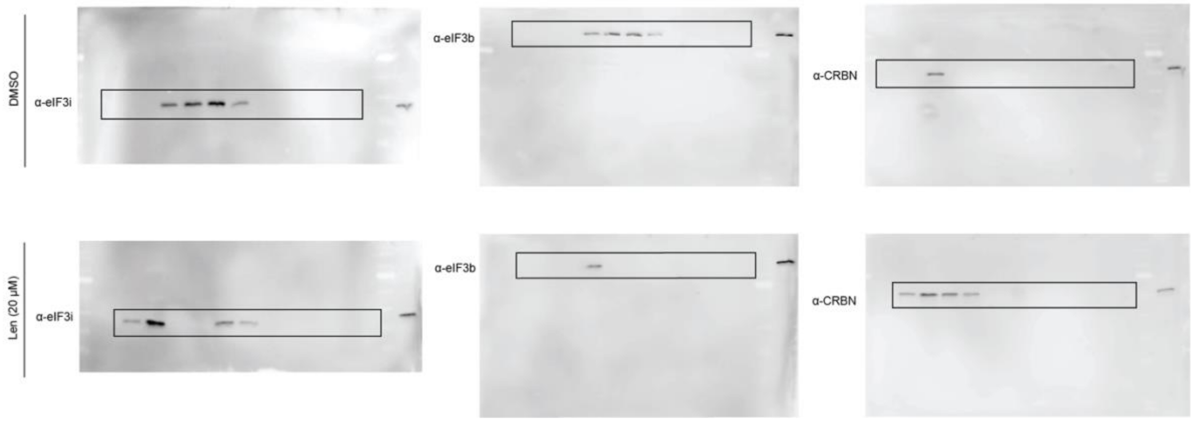


**Figure S14 |** Uncropped immunoblots related to **Figure 1.**

**Figure 2d**

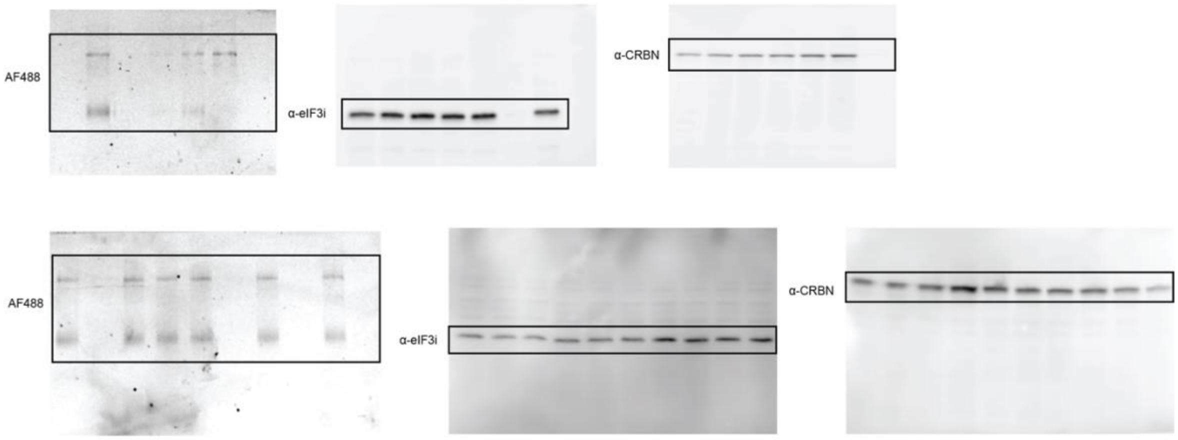


**Figure 2e**

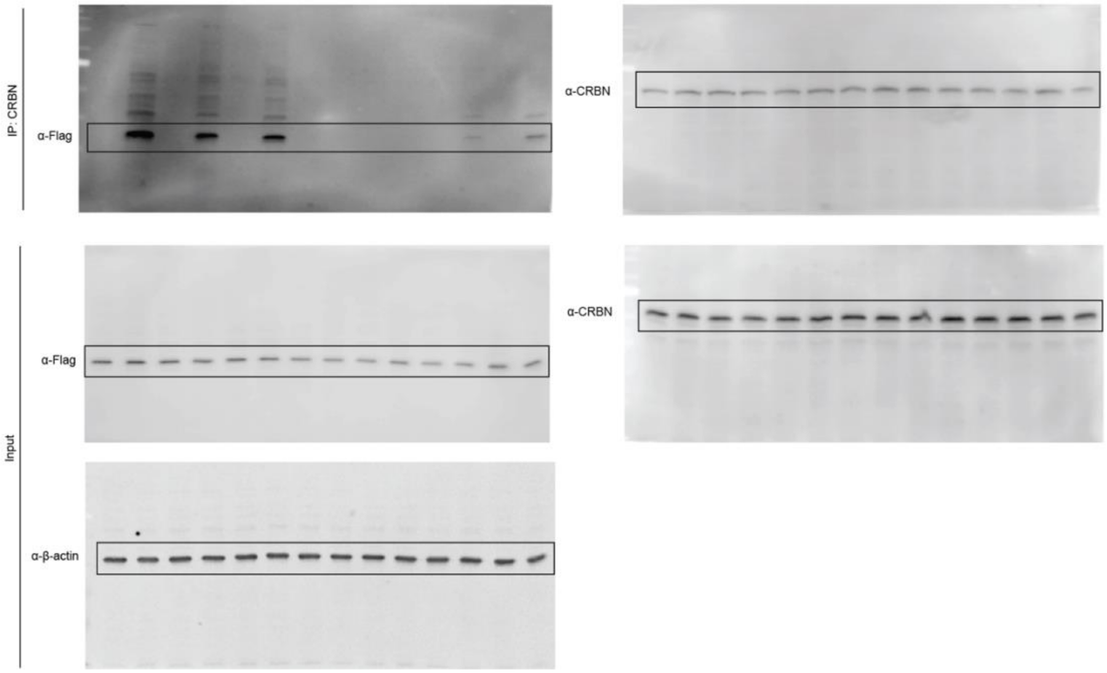


**Figure S15 |** Uncropped immunoblots related to **Figure 2.**

**Figure 3b**

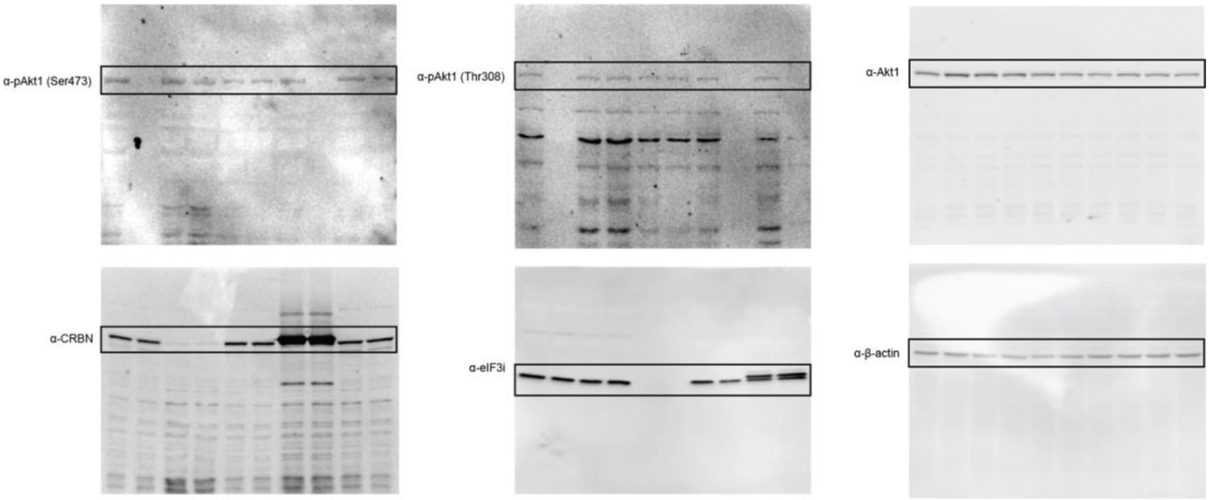


**Figure 5b**

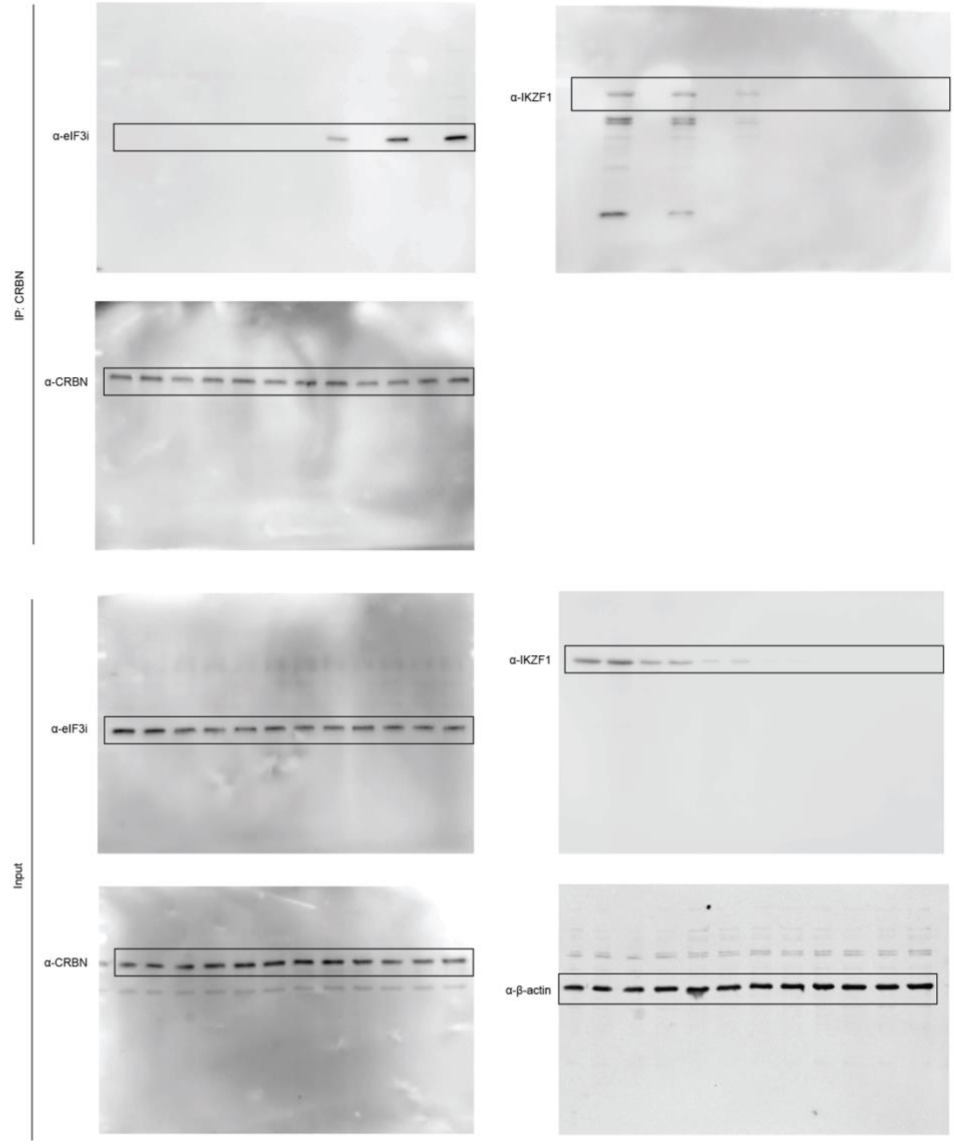


**Figure S16 |** Uncropped immunoblots related to **Figure 3** and **Figure 5.**

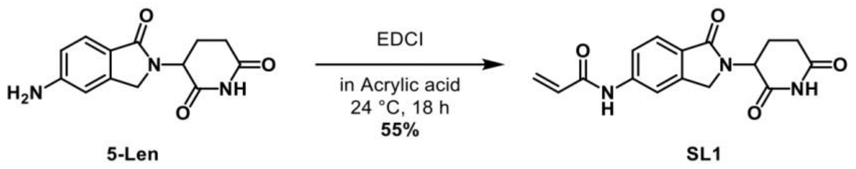


**Scheme S1 |** Synthetic scheme for SL1.

### Methods

#### General methods and instrumentation.

Photoirradiation was performed using a Dymax ECE 5000 UV Light-Curing Flood Lamp system (Cat # 41060) in a 4 ºC cold room unless otherwise noted. The lamp was turned on for at least 15 minutes prior to use.

Protein quantification was performed by bicinchoninic acid assay on a multimode microplate reader FilterMax F3 (Molecular Devices LLC, Sunnyvale, CA). Sonication of cells or protein pellets was performed using a Branson SFX250 Sonifier (Branson Ultrasonics, Brookfield, CT). In-gel fluorescence were detected on an Azure Imager C600 (Azure Biosystems, Inc., Dublin, CA). AlphaScreen assay was performed on a multimode microplate reader SpectraMax i3x. DNA or RNA concentration was measured with a NanoDrop One UV-vis spectrophotometer (Thermo Fisher, Cat # ND-ONE-W).

Proteomics experiments were performed on a Lumos Tribrid Orbitrap (Thermo Scientific) equipped with an UltiMate 3000 RSLCnano System (Thermo Scientific) within the Mass Spectrometry and Proteomics Resource Laboratory at Harvard University (MSPRL). Peptides were dried using an Eppendorf Vacufuge (Speed-Vac). Cell sorting experiments were performed on a BD FACSAria II cell sorter (BD Bioscience). Flow cytometry quantification were performed on BD LSR Fortessa (BD Bioscience, SORP) and BD LSR II flow cytometric analyzers (BD Biosciences, SORP).

For immunoblotting analysis, proteins were transferred from SDS-PAGE gels to nitrocellulose membranes using iBlot-2 dry blotting system (Thermo Scientific). Membranes were blocked with Tris buffered saline containing 0.1% Tween-20 and 5% BSA and incubated with the primary antibodies and the secondary antibodies sequentially. Immunoblots images were captured by Azure Imager C600 (Azure Biosystems, Dublin, CA).

#### General chemical methods and instrumentation.

DMSO-$d_6$ was purchased from Cambridge Isotope Laboratories. All other solvents and reagents were purchased from chemical suppliers (Sigma Aldrich, Wuxi AppTech, Acros Organics, Thermo Fisher Scientific or VWR Chemicals BDH®) and were used as received unless otherwise noted. Chemicals purchased from BioVision include lenalidomide (Cat # 1862-25), thalidomide (Cat # 10188-734), CC-220 (Cat # B2284-1) and dBET6 (Cat # 76411-760). Photolenalidomide was prepared according to Woo and co-workers.[1] Cycloheximide (CHX) was purchased from Sigma-Aldrich (Cat # C4859). Chemicals purchased from Selleck Chemicals include CC-885 (Cat # S8300) and CC-90009 (Cat # S9832).

Flash Column Chromatography was performed using silica gel purchased from Silicycle (SilicaFlash® F60, 40–63 µm) with the aid of a CombiFlash® NextGen 300+ Automated Flash Chromatography System (Teledyne ISCO). Organic solutions were concentrated in vacuo using a Buchi rotary evaporator. Analytical thin-layer chromatography (TLC) was performed using glass plates pre-coated with silica gel (0.25 mm, 60 Å pore size) embedded with a fluorescent indicator (254 nm). TLC plates were visualized by exposure to ultraviolet light (UV) or iodine ($I_2$).

Proton nuclear magnetic resonance spectrum ($^1$H NMR) and carbon nuclear magnetic resonance spectrum ($^{13}$C NMR) were recorded on a Bruker 400 MHz instrument at 25 °C. Chemical shifts were reported in parts per million (ppm, δ scale) relative to residual solvent as an internal reference (DMSO: 2.50 ppm for $^1$H and 39.52 ppm for $^{13}$C). Data are represented as follows: chemical shift, multiplicity (s = singlet, d = doublet, t = triplet, q = quartet, quin = quintet, m = multiplet and/or multiple resonances, br = broad, app = apparent), integration, coupling constant in Hertz, and assignment. Infrared (IR) spectrum was recorded on a Bruker

ALPHA FT-IR and are reported in terms of frequency of absorption ($cm^{-1}$) and intensity of absorption (s = strong, m = medium, w = weak, br = broad). High-resolution mass spectra were recorded on a Bruker microTOF-Q II hybrid quadrupole-time of flight system and on an Agilent 1260 UPLC-MS system.

**Synthetic procedures.**

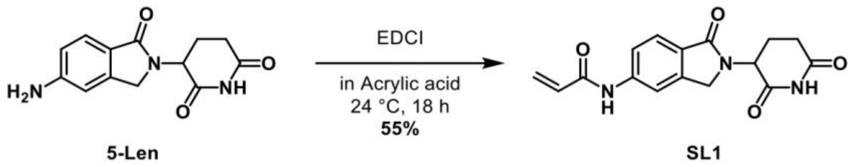


**Synthesis of *N*-(2-(2,6-dioxopiperidin-3-yl)-1-oxoisoindolin-5-yl)acrylamide (SL1):**

5-$NH_2$-Lenalidomide (30.0 mg, 116 μmol, 1.0 equiv.) was added in a single portion to a stirred solution of acrylic acid (500 μL, 63 equiv.) in a 20 mL vial. EDCI (26.7 mg, 139 μmol, 1.20 equiv.) was added into the mixture. The reaction mixture was stirred for 16 h at 24 ℃. TLC showed full conversion of the starting material. The product mixture was then loaded on silica gel directly and subjected to purification by flash column chromatography (ISCO 12 g gold column, 0 to 10% gradient MeOH in DCM over 20 min) to afford the product **SL1** as a white solid (20.0 mg, 63.9 μmol, 55%).

**$^{1}$H NMR** (400 MHz, DMSO) δ 10.98 (s, 1H), 10.47 (s, 1H), 8.05 (t, *J* = 1.2 Hz, 1H), 7.74 – 7.55 (m, 2H), 6.48 (dd, *J* = 17.0, 10.1 Hz, 1H), 6.31 (dd, *J* = 17.0, 2.0 Hz, 1H), 5.81 (dd, *J* = 10.1, 2.0 Hz, 1H), 5.09 (dd, *J* = 13.3, 5.1 Hz, 1H), 4.45 (d, *J* = 17.5 Hz, 1H), 4.31 (d, *J* = 17.2 Hz, 1H), 2.91 (ddd, *J* = 17.2, 13.6, 5.4 Hz, 1H), 2.64 – 2.55 (m, 1H), 2.43 – 2.31 (m, 1H), 1.99 (dtd, *J* = 12.6, 5.2, 2.2 Hz, 1H). **$^{13}$C NMR** (101 MHz, DMSO) δ 172.9, 171.1, 167.8, 163.5, 143.3, 142.2, 131.6, 127.7, 126.6, 123.7, 119.0, 113.5, 51.5, 47.1, 31.2, 22.5. **IR** (ATR-FTIR), in DMSO, $cm^{-1}$: 3487 (br), 2996 (w), 2912 (w), 1686 (w), 1436 (m), 1407 (m), 1310 (m), 1235 (w), 1204(w), 1042 (s), 1020 (s), 952 (s), 930 (m), 897 (w). **HRMS-ESI** (*m/z*): $[M+H]^+$ calculated for $C_{16}H_{16}N_3O_4$ 314.1135; found, 314.1135.

**Antibodies and biological reagents.**

Antibodies included anti-eIF3i (Proteintech, Cat # 10086-380), anti-CRBN (for IP: Cell Signaling Technology, Cat # 71810S, for WB: Sigma-Aldrich, Cat # HPA045910), anti-eIF3b (Bethyl Laboratories, Cat # A301-760A-T), anti-eIF3g (Bethyl Laboratories, Cat # A301-757A-T), anti-pAkt1 (Ser473, Cell Signaling Technology, Cat # 4060), anti-pAkt1 (Thr308, Cell Signaling Technology, Cat # 13038), anti-Akt1 (Cell Signaling Technology, Cat # 2938), anti-IKZF1 (Cell Signaling Technology, Cat # 9034S), anti-IKZF3 (Novus Bio, Cat # NBP2-16938), anti-Flag (Sigma-Aldrich, Cat # F3165), anti-β-actin (Santa Cruz, Cat # sc-47778),

Antibody-conjugated beads for immunoprecipitation included protein A magnetic beads (Cell Signaling Technology, Cat # 73778), protein G magnetic beads (Cell Signaling Technology, Cat # 70024), Anti-Flag M2 magnetic beads (Sigma-Aldrich, Cat # M8823), anti-EPEA CaptureSelect™ C-tag affinity matrix (Thermo Scientific, Cat # 191307005), and Pierce Anti-HA magnetic beads (Pierce, Cat # PI88836). For AlphaScreen assay, nickel chelate acceptor beads (PerkinElmer, Cat # AS101D) and glutathione donor beads (PerkinElmer, Cat #6765300) were used. Cell lysis buffer was prepared by diluting from 10X cell lysis buffer (Cell Signaling Technology, Cat # 9803S). TBS buffer was prepared by diluting from 10X TBS buffer solution (Quality Biological, Cat # 10128-546). For qRT-PCR, RNeasy Plus Mini kit (Qiagen, Cat # 74134), PrimeScript RT reagent kit (TakaraBio Cat # RR037A), and QuantiTect SYBR Green PCR kit (Qiagen, Cat # 204143) were used. For Akt1 phosphorylation ELISA assay, FastScan™ Phospho-Akt (Ser473) ELISA kit (Cell Signaling Technology, Cat # 80895C) was used. Sequencing-grade modified trypsin (Promega, Cat # V5111), sequencing-grade chymotrypsin (Promega, Cat # V1061) and mini Bio-Spin columns (Bio-Rad, Cat # 7326207) were purchased.

**Cell culture and transfection.**

HEK293T cells (a gift from the Bertozzi lab, Stanford University), HEK293T cells stably expressing Flag-CRBN (HEK293T-CRBN) and HEK293T cells with CRBN knock-down by lentiviral shRNA (a gift from the Deshaies lab, California Institute of Technology), A549 (ATCC), and RAW264.7 cells (a gift from the Mitchison lab, Harvard Medical School) were cultured in Dulbecco's Modified Eagle Medium (DMEM, Cat # 11995073), supplemented with penicillin (50 μg/mL), streptomycin (50 μg/mL), and 10% (v/v) FBS. HEK293T-CRBN cell line was established by transducing cells with the corresponding lentiviral particles with transduction efficiency monitored by GFP signals via flow cytometry. MM.1S cells (ATCC), U937 cells (ATCC), Jurkat cells (ATCC) and MOLM13 cells (a gift from the Shair lab, Harvard University) were cultured in RPMI1640 (Cat #45000-396) supplemented with penicillin (50 μg/mL), streptomycin (50 μg/mL) and 10% (v/v) FBS. HUVECs (Lonza) were cultured in endothelial cell growth basal medium EBM-2 (Cat # CC-3156) supplemented with Bovine Brain Extract, rhEGF, L-glutamine, heparin sulfate, hydrocortisone hemisuccinate, Fetal Bovine Serum, and ascorbic acid from the endothelial cell growth EGM-2 kit (Cat # CC-3162) as instructed. For in vitro angiogenesis assays, only HUVECs between passage 4-8 was used according to published protocols.[2-3]

Transfection was performed using TransIT-PRO (Mirus Bio, Cat # MIR 5740) according to the manufacturer's instructions. Cells were lysed with 1X cell lysis buffer diluted from 10X cell lysis buffer supplemented with 1X cOmplete protease inhibitor cocktail (PI, Cat # 11697498001) unless otherwise noted.

**Plasmids and subcloning.**

Human CRBN with a Flag tag, human eIF3i with a His tag and human eIF3i with a Flag tag were obtained as detailed in the table below. Mutant plasmids of eIF3i or CRBN were constructed by subcloning. Empty pcDNA3.1 vector was used as a negative control for transfection.

Genetic constructs and primers used in this study are listed below:

| ***No.*** | ***Plasmid name*** | ***Abbreviated name*** | ***Details*** | ***Source*** |
|---|---|---|---|---|
| 1 | pcDNA3.1 | Ctrl | Empty vector for mammalian expression | Invitrogen, Cat # V79020 |
| 2 | pCDH-Flag-CRBN-WT-T2AcGFP-MSCV | Flag-CRBN (stable transfection) | Full length CRBN with a Flag tag on pCDH vector | A gift from Deshaies lab |
| 3 | pcDNA3-FLAG-CRBN-pGK-HYG | Flag-CRBN (transient transfection) | Full length CRBN with a Flag tag on pcDNA vector | Addgene, Cat # 107380 |
| 4 | pcDNA3-FLAG-CRBN (YWAA)-pGK-HYG | Flag-CRBN (YWAA mutant) | Full length CRBN with mutations Y384A, W386A and a Flag tag on pcDNA vector | Made in house |
| 5 | pCMV3-N-His-eIF3i | His-eIF3i | Full length eIF3i with a His tag on pCMV vector | SinoBio, Cat # HG6651 |
| 6 | pcDNA3.1+eIF3i-C-Flag | eIF3i-Flag (WT) | Full length eIF3i with a Flag tag on pcDNA vector | GenScript, Cat # OHu12097D |
| 7 | pcDNA3.1+eIF3i(C120A)-C-Flag | eIF3i-Flag (C120A) | Full length eIF3i with mutation C120A and Flag tag on pcDNA vector | Made in house |
| 8 | pcDNA3.1+eIF3i(C57A)-C-Flag | eIF3i-Flag (C57A) | Full length eIF3i with mutation C157A and Flag tag on pcDNA vector | Made in house |
| 9 | pcDNA3.1+eIF3i(C76A)-C-Flag | eIF3i-Flag (C76A) | Full length eIF3i with mutation C76A and Flag tag on pcDNA vector | Made in house |
| 10 | pcDNA3.1+eIF3i(C160A)-C-Flag | eIF3i-Flag (C160A) | Full length eIF3i with mutation C160A and Flag tag on pcDNA vector | Made in house |
| 11 | pcDNA3.1+eIF3i(C120Y)-C-Flag | eIF3i-Flag (C120Y) | Full length eIF3i with mutation C120Y and Flag tag on pcDNA vector | Made in house |
| 12 | pcDNA3.1+eIF3i(Q119W)-C-Flag | eIF3i-Flag (Q119W) | Full length eIF3i with mutation A119W and Flag tag on pcDNA vector | Made in house |
| 13 | pcDNA3.1+eIF3i(Y118W)-C-Flag | eIF3i-Flag (Y118W) | Full length eIF3i with mutation Y118W and Flag tag on pcDNA vector | Made in house |
| 14 | pcDNA3.1+eIF3i(G117W)-C-Flag | eIF3i-Flag (G117W) | Full length eIF3i with mutation G117W and Flag tag on pcDNA vector | Made in house |
| 15 | pcDNA3.1+eIF3i(M116W)-C-Flag | eIF3i-Flag (M116W) | Full length eIF3i with mutation M116W and Flag tag on pcDNA vector | Made in house |
| 16 | pcDNA3.1+eIF3i(Q115W)-C-Flag | eIF3i-Flag (Q115W) | Full length eIF3i with mutation Q115W and Flag tag on pcDNA vector | Made in house |

**Immunoprecipitation assays in live cells.**

For immunoprecipitation of proteins with the Flag tag, cell lysates with equal amounts of protein were diluted with the lysis buffer and incubated with Anti-Flag M2 magnetic beads (pre-washed three times with TBS) for 2 h at 4 ˚C with rotation. The beads were washed three times with TBS buffer (50 mM Tris HCl, 150 mM NaCl, pH 7.4). The enriched proteins were eluted with SDS sample buffer for Western blot analysis or 50 µL 3X Flag peptide solution (final concentration 150 ng/µl), according to the manufacturer's instructions.

For immunoprecipitation of proteins with the His tag, cell lysates with equal amounts of protein were diluted with PBS and incubated with C-tag affinity matrix (pre-washed three times with PBS) for 2 h at 25 ˚C. The beads were washed three times with PBS buffer. The enriched proteins were eluted with SDS sample buffer for Western blot analysis or 50 µL neutral pH-based elution buffer (20 mM Tris, 2 M $MgCl_2$, pH 7.0), according to the manufacturer's instructions.

For endogenous protein immunoprecipitation using protein A/G beads, cell lysates with equal amounts of protein were diluted with PBS and incubated with protein A/G beads (pre-washed three times with binding buffer, 50 mM Tris, 150 mM NaCl, 0.2% Triton, pH=7.5) for 2 h at 4 ˚C along with the protein-specific antibody at the vendor-suggested dilution. The beads were washed three times with binding buffer (50 mM Tris, 150 mM NaCl, 0.2% Triton, pH=7.5). The enriched proteins were eluted with acidic elution buffer (100 mM glycine, 0.1% Triton, pH=2.8) before neutralizing with 1M Tris (pH=8), according to the manufacturer's instructions.

**Sucrose gradient centrifuge fractionation assay.**

Sucrose gradient separation was performed and adapted according to the method from Moazed and co-workers.[4] In brief, sucrose gradients (10%-50%) were prepared using the Gradient station (BioComp, Cat # B153-002). An Optima TPX Ultracentrifuge (Beckman Coulter) equipped with SWI-44 rotor was used for ultracentrifugation for indicated time at 4˚C with 35k rpm. Gradients of 2.2 mL were fractionated into fractions by pipetting from top. The separated fractions were then analyzed by Western blot individually or combined before Western blot analysis.

**AlphaScreen assay for binding interaction analysis.**

AlphaScreen buffer was freshly made containing 50 mM HEPES 200 mM NaCl, 0.1% w/v BSA, 1 mM TCEP (pH=7.4) right before the assay. Compounds (lenalidomide, photolenalidomide or SL1) at the concentrations indicated were mixed in AlphaScreen buffer along with indicated proteins at their respective concentrations (His-CRBN/DDB1 50 nM, GST-eIF3i 100 nM), and 10 µL of the reaction mixture was incubated at 25 ˚C for 1 h in a 384-well white opaque OptiPlate384 plate. Following the incubation, 5 µL of the detection mixture containing nickel chelate acceptor beads (PerkinElmer, Cat # AS101D) and glutathione donor beads (PerkinElmer, Cat #6765300) were added to each well in dark. After incubation at 25 ˚C for 1 h in dark, luminescent signals were detected using a multimode microplate reader SpectraMax i3x.

**Direct covalent labeling with SL1 and MS sample preparation.**

Direct covalent labeling was performed according to the protocol by Nomura and co-workers.[5] Purified CRBN (2 µg) and eIF3i (2 µg) in 50 µL 1% Triton in PBS was incubated at 25 ˚C for 30 min with 50 µM SL1. Then 200 µL pre-chilled acetone was added and the proteins were precipitated and separated by centrifugation (4℃, 10 min, 21,300 x g). The supernatant was discarded, and the protein pellet was air dried for 10 min at 25℃. The protein pellet was resuspended in 50 µL freshly made 50 mM TEAB with 8 M urea by vortex. The protein was then diluted in 850 µL 50 mM TEAB to a final concentration of 0.5 M urea and digested by sequence-grade trypsin (0.5 µg) for 18 h at 37℃ with inversion. The digested protein was then dried on a Speed-Vac to dryness before desalting using a Pierce C18 tip according to the manufacture's protocol. The desalted peptides were dried on a Speed-Vac before they were re-suspended in 20 µL 0.1% formic acid/water for analysis.

**Mass spectrometry sample processing for covalent binding site identification.**

The sample was loaded onto a microcapillary trapping column (C18 Reprosil resin, Dr. Maisch, 5 µm particle size, 100 Å pore size, 30 mm length, 100 µm internal diameter) and then separated on an analytical column (50 cm µPAC™ column, PharmaFluidics) at 200 nL/min. The column temperature was maintained at 40℃. Peptides were eluted with a water/acetonitrile gradient (buffer A = 0.1% formic acid/water, buffer B = 0.1% formic acid/acetonitrile; flow rate 200 nL/min; gradient: hold at 2% B for 5 min, increase to 6% B over 1 min, increase to 35% B over 29 min, increase to 95% B over 5 min, hold at 95% B for 20 min). Electrospray ionization was enabled through applying a voltage of 1.8 kV using a home-made electrode junction at the end of the microcapillary column and sprayed from stainless-steel needle (PepSep, Denmark). The Lumos Orbitrap was operated in data-dependent mode and survey scans of peptide precursors were performed at 120K FWHM resolution over a *m/z* range of 400-1800, an AGC setting of 1,000,000, and the maximum ion accumulation time of 50 ms. Tandem MS was performed on the 10 most abundant precursors with a standard AGC setting and collision energy of 35% for CID, an AGC setting of 200% and fragmentation energy of 37% for HCD together with an AGC setting of 600% and SA energy of 37% for EThcD. An isolation window of 3 *m/z* used to ensure capture of isotopically coded species. With a mass tolerance of 10 ppm, selected precursors were excluded from further fragmentation for 60 s.

**Mass spectrometry data analysis procedure for covalent binding site identification**.

The raw data were analyzed using Proteome Discoverer 2.4. Assignment of MS/MS spectra was performed using the Sequest HT algorithm by searching the data against CRBN, eIF3i and common contaminant proteins. Search parameters included: mass tolerance of 10 ppm for the precursor, 0.02 Da for HCD fragment ions, 0.6 Da for CID fragment ions, semi-specific trypsin digestion, 2 missed cleavages, and a dynamic modification of the compounds on any amino acid residues (SL1, +313.1063). Peptide spectral matches (PSMs) were validated with the Target Decoy PSM Validator. Spectra assigned as probe-conjugated peptides were manually validated by evaluating the isotopic coding embedded in the MS1 precursor.

**Covalent competitive displacement PAL assay.**

Covalent competitive displacement assay was adapted from photoaffinity labeling and in-gel fluorescence assay from Woo and co-workers.[1] Purified Flag-CRBN (0.5 µg) and/or eIF3i-Flag (0.5 µg) in 50 µL 1% Triton in PBS was pre-incubated with 50 µM SL1 at 25 ˚C for 30 min. Indicated concentration of photolenalidomide was then added and incubated drugs for 1 h at 37 ˚C before photoirradiation for 60 s at 4 ˚C via Dymax. The proteins were then precipitated by the addition of 200 µL pre-chilled acetone and centrifugation (4℃, 10 min, 21,300 x g). After separation, the protein pellets were resuspended in PBS and then tagged with AlexaFluor® 488 azide via CuAAC by addition of a pre-mixed cocktail (final concentrations: 25 µM AlexaFluor® 488 azide, 100 µM THPTA, 1 mM $CuSO_4$, 2 mM freshly dissolved sodium ascorbate) and incubated for 1 h at 25 ˚C with inversion. The reactions were then quenched by SDS sample buffer before Western blot analysis.

**Structural molecular modeling.**

Molecular docking study was performed using Molecular Operating Environment (MOE) version 2020.09.[6] CRBN was extracted from DDB1-CRBN-lenalidomide structure (PDB:4CI2) along with ligand lenalidomide. eIF3i was extracted from human 48S translational initiation complex eIF3b:g:i structure (PDB: 6YBT) and compared with eIF3i structure from AlphaFold prediction (AlphaFold-Q13347-F1-model_v2, https://alphafold.ebi.ac.uk/entry/Q13347). The input coordinates of both structures were processed first using the protein preparation function by adding hydrogen atoms, energy minimization with Amber10:EHT force field and protonation by Protonate 3D. The complex model was obtained using the protein-protein dock tool in MOE, where the docking algorithm starts with a coarse-grained model, and then generates a set of uniformly distributed rotations by Hopf fibration. Initial docked poses were generated and filtered before they were subjected to coarse refinement. The output database consisted of 100 poses which were generated by triangle match placement method with the side chain position refined by induced fit. The poses were scored by the estimation of the free energy of the complex (London ΔG) and rescored with GVI/WSA function.

**Polysome profiling.**

Polysome profiling assay was performed according to Chassé, Boulben, Morales and co-workers.[7] Cells after treatment were harvest and lysed. Cell lysates were then separated by sucrose gradient separation before each fraction was collected. The concentration of nucleic acid in each fraction was measured by absorbance at A260 nm by NanoDrop One.

**RNA isolation and qRT-PCR experiment.**

Cells were treated as indicated before RNA isolation. Qiagen RNeasy-Plus Mini kits were used to purify RNA from cells, according to the instructions from the manufacturer. Isolated RNA then underwent two-step real-time RT-PCR by using PrimeScript RT reagent kit for reverse transcription and QuantiTect SYBR Green PCR kit.

**Akt1 phosphorylation ELISA assay.**

Akt1 phosphorylation was quantified using ELISA kit (Cell Signaling Technology, Cat # 80895) according to the manufacturers' instructions. In brief, cells treated with indicated compounds were harvested and lysed on ice with pre-chilled cell lysis buffer (Cell Signaling Technology) before the cell lysates was cleared by centrifugation (4 ℃, 10 min, 21,300 x g). The lysates (50 µL) were then added to the FastScan$^{TM}$ 96-well ELISA strip plate before 50 µL antibody cocktail was added. The plate was then sealed and incubated for 1 h at 25 ˚C with moderate agitation. After incubation, the wells were washed with 200 µL wash buffer (freshly prepared by diluting 20X ELISA Wash Buffer provided in the kit) for three times before 100 µL TMB substrate (provided in the kit) was added. The plate was then incubated for 10 min at 37 ˚C before 100 µL stop solution (provided in the kit) was added to each well with moderate agitation for 2 min at 25 ˚C. Absorbance at 450 nm was then detected using SpectraMax i3x plate reader.

**Wound healing assay.**

HUVEC cells were plated in complete EGM-2 media on a rectangular cell culture plate (Cat # CS210543). The confluent monolayer of HUVEC cells were then wounded by using the Cell Comb™ Scratch Assay Kit (MilliporeSigma, Cat # 1710191) and washed three times with HBSS buffer. The remaining cells were treated with the indicated compound in complete EGM-2 media for 24 h at 37 ˚C before they were stained with Calcein AM for imaging.

**Transwell migration assay with Boyden chambers.**

HUVEC cells were harvested and resuspended with EGM-2 basal medium and added to the top chamber of the transwell inserts (Celltreat, 3.0 µm pore size, Cat # 270-10043-ER). Bottom chambers were filled with complete EGM-2 medium with DMSO or compound treatment. Cells were allowed to migrate for 24 h at 37 ˚C. Noninvasive cells were removed by cotton swab scrubbing and invasive cells were fixed with 4% formaldehyde and methanol before they were stained with DAPI solution for imaging.

**Tube formation assay.**

Matrigel (Corning, Cat # 356237) was thawed for 16 h at 4 ˚C before addition to each well of a chilled 48-well glass bottom plate with chilled pipette tips and was allowed to polymerize for 30 min at 37 ˚C. HUVEC cells were plated on top of the polymerized Matrigel and treated with the indicated compound in complete EGM-2 media for 24 h at 37 ˚C before they were stained with Calcein AM for imaging.

**Quantitative mass spectrometry sample preparation.**

Cells were treated with indicated compounds and lysed before the cell lysate was subjected to coimmunoprecipitation. Eluted proteins were then digested by sequence-grade trypsin (1 µg) for 18 h at 37 ℃ with inversion. The digested protein was then dried on a Speed-Vac to dryness before desalting using a Pierce C18 tip according to the manufacture's protocol. Desalted peptides were then resuspended in 50 µL of 50 mM TEAB buffer (pH=8.5), treated with 5 µL of the TMT 16-plex reagents (11.9 µg/µL) and then incubated for 1 h at 25 ˚C. The TMT labeling was quenched with 2 µL of 5% hydroxylamine solution for 15 min at 25 ˚C. The TMT-labeled peptides were then combined and dried on the Speed-Vac. The peptides were resuspended in 100 µL 1% formic acid/water and desalted with C18 Tips (Zip-Tip) according to the manufacturer's instructions and stored at -20 ˚C until analysis.

**High pH separation and quantitative mass spectrometry analysis.**

The sample was separated using the Pierce™ High pH Reversed-Phase Peptide Fractionation Kit (Thermo Fischer) according to manufacturer's instructions. After separation, each fraction was submitted for a single LC-MS/MS experiment that was performed on a Lumos Tribrid Orbitrap equipped with an UltiMate 3000 RSLCnano System. Peptides were separated onto a 100 µm inner diameter microcapillary trapping column packed first with approximately 3 cm of C18 Reprosil resin (5 µm, 100 Å, Dr. Maisch GmbH, Germany) followed by analytical column 50 cm microcapillary based PharmaFluidics (Belgium). Separation was achieved through applying a gradient from 5–27% acetonitrile in 0.1% formic acid in water over 90 min at 200 nl $min^{-1}$. Electrospray ionization was enabled through applying a voltage of 1.8 kV using a home-made electrode junction at the end of the microcapillary column and sprayed from stainless-steel needle (PepSep, Denmark). The Lumos Orbitrap was operated in data-dependent mode for the mass spectrometry methods. The mass spectrometry survey scan was performed in the Orbitrap in the range of 400–1,800 m/z at a resolution of $1.2 \times 10^5$, followed by the selection of the 10 most intense ions (TOP10) for CID-MS2 fragmentation in the Ion trap using a precursor isolation width window of 2 m/z, an AGC setting of 10,000, and a maximum ion accumulation of 200 ms. Singly charged ion species were not subjected to CID fragmentation. Normalized collision energy was set to 35 V and an activation time of 10 ms. Ions in a 10-ppm m/z window around ions selected for MS2 were excluded from further selection for fragmentation for 60 s. The same TOP10 ions were subjected to HCD MS2 event in Orbitrap part of the instrument. The fragment ion isolation width was set to 0.8 Da, with 0.3 Da offset, AGC was set to 50,000, the maximum ion time was 100 ms, normalized collision energy was set to 34 V and an activation time of 1 ms for each HCD MS2 scan.

**Quantitative mass spectrometry data analysis**.

The raw data were analyzed using Proteome Discoverer 2.4.1.15. Assignment of MS/MS spectra was performed using the Sequest HT algorithm by searching the data against a protein sequence database including all entries from the Human Uniprot database (Aug 19, 2016, 20,156 total entries, SwissProt) and a list of common contaminant proteins. Search parameters included: a 10-ppm precursor ion tolerance and 0.02 Da fragment ion tolerance for HCD or 0.6 Da fragment ion tolerance for CID, full tryptic protease specificity with up to two missed cleavages, a static modification of 16-plex TMTpro tags on peptide N-termini and lysine residues (+304.2071 Da), a static modification of cysteine carbamidomethylation (+57.0214 Da), a dynamic modification of methionine oxidation (+15.9949 Da), a dynamic modification of deamidation on asparagine and glutamine residues (+0.9847 Da) and a dynamic modification of acetylation on peptide N-termini (+42.0106 Da). Peptide spectral matches (PSMs) were filtered for a false discovery rate (FDR) of 1% following a target-decoy analysis with Percolator. Reporter ions were quantified using a 0.02 m/z window centered on the theoretical m/z value and the intensity of the signal closest to the theoretical m/z value was recorded. Reporter ion intensities were exported in result file of Proteome Discoverer search engine as an excel tables. The total signal intensity across all peptides quantified was summed and normalized for each TMT channel. Quantified proteins were required to have at least three unique peptides and four separate PSMs. For P-value and fold change calculations, the data were further processed using a custom algorithm as described in previous protocol.[8] All TMT-based experiments were performed with four replicates.

**Software.**

Data were analyzed and visualized using Microsoft Excel (v16.22) and GraphPad Prism (v8.0.1), in addition to software listed by experiment. NMR data were analyzed using MestReNova (v14.0.1). DNA and protein sequences were analyzed using Geneious (v10.0.7). FACS data were analyzed by FlowJo (v10.6.1). Proteomics data were analyzed by Proteome Discoverer (v2.4.1.15). Images were made using ImageJ (NIH, v1.50i), ImageStudioLite (v5.2.5), and Adobe Illustrator (v22.1). Docking studies were performed with Molecular Operating Environment (MOE, v2020.09) and the obtained structures were analyzed with PyMOL (v2.3.1).

**Statistical analysis.**

Statistical analyses (unpaired Student's *t*-tests) were performed using GraphPad Prism. Data were derived from at least three biological replicate experiments and presented as the mean ± s.d., *P< 0.0332, **P< 0.0021, ***P < 0.0002, ****P < 0.0001 and n.s., not significant.

**NMR Spectra of SL1**

$^{1}$H-NMR of **SL1**

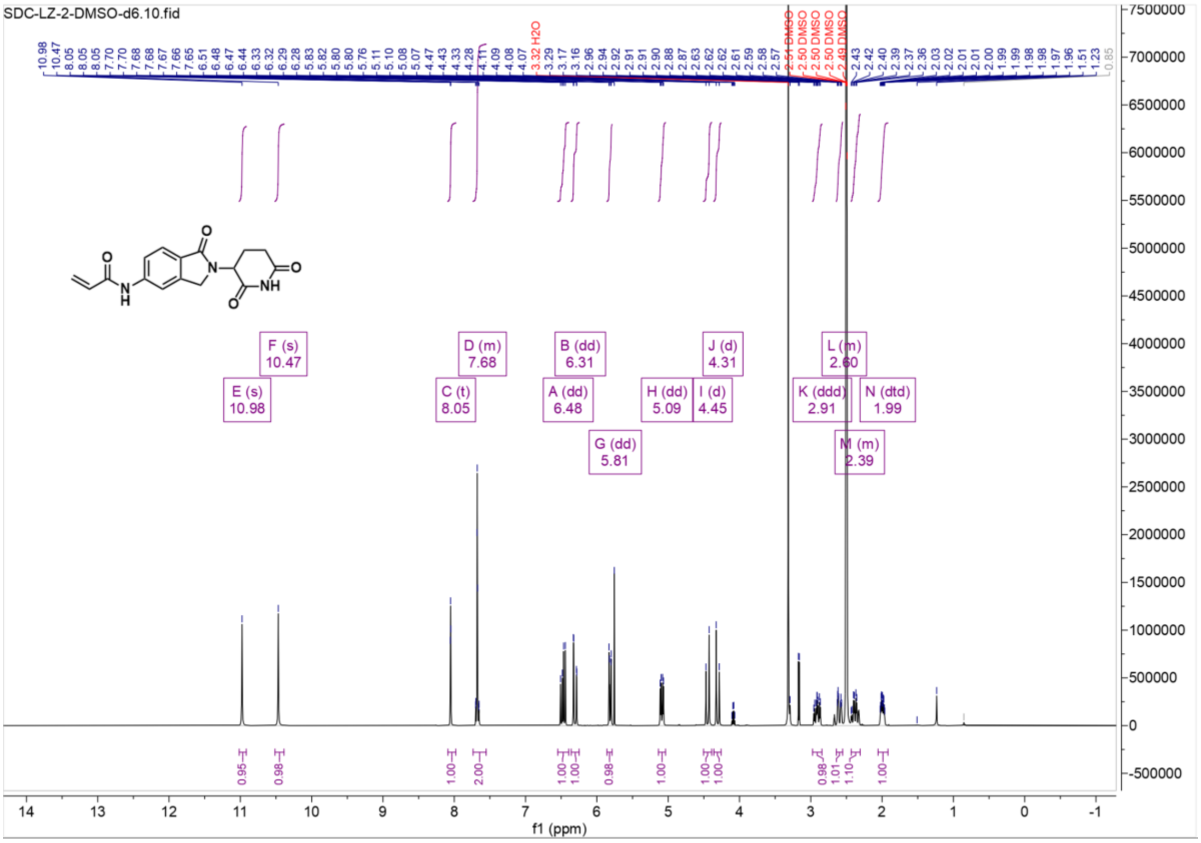

$^{13}C$-NMR of **SL1**

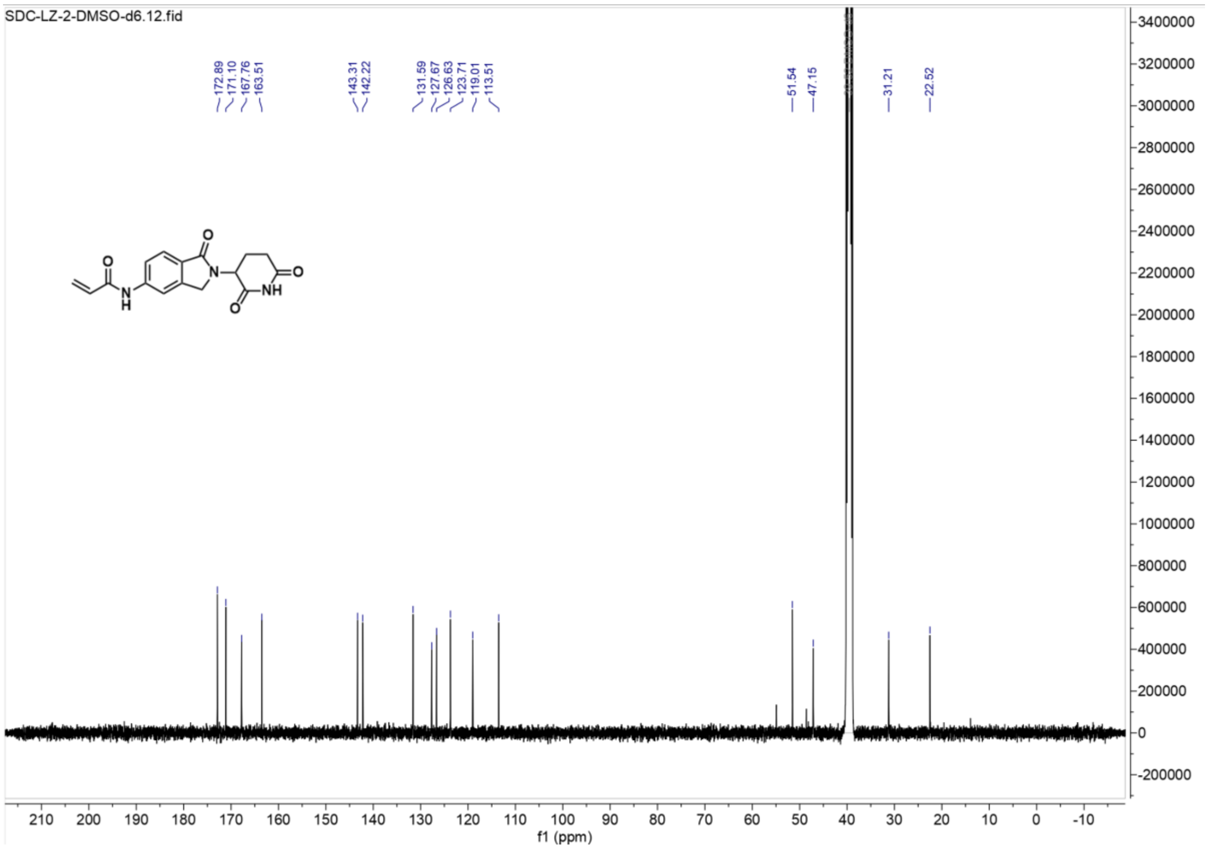

IR of **SL1**:

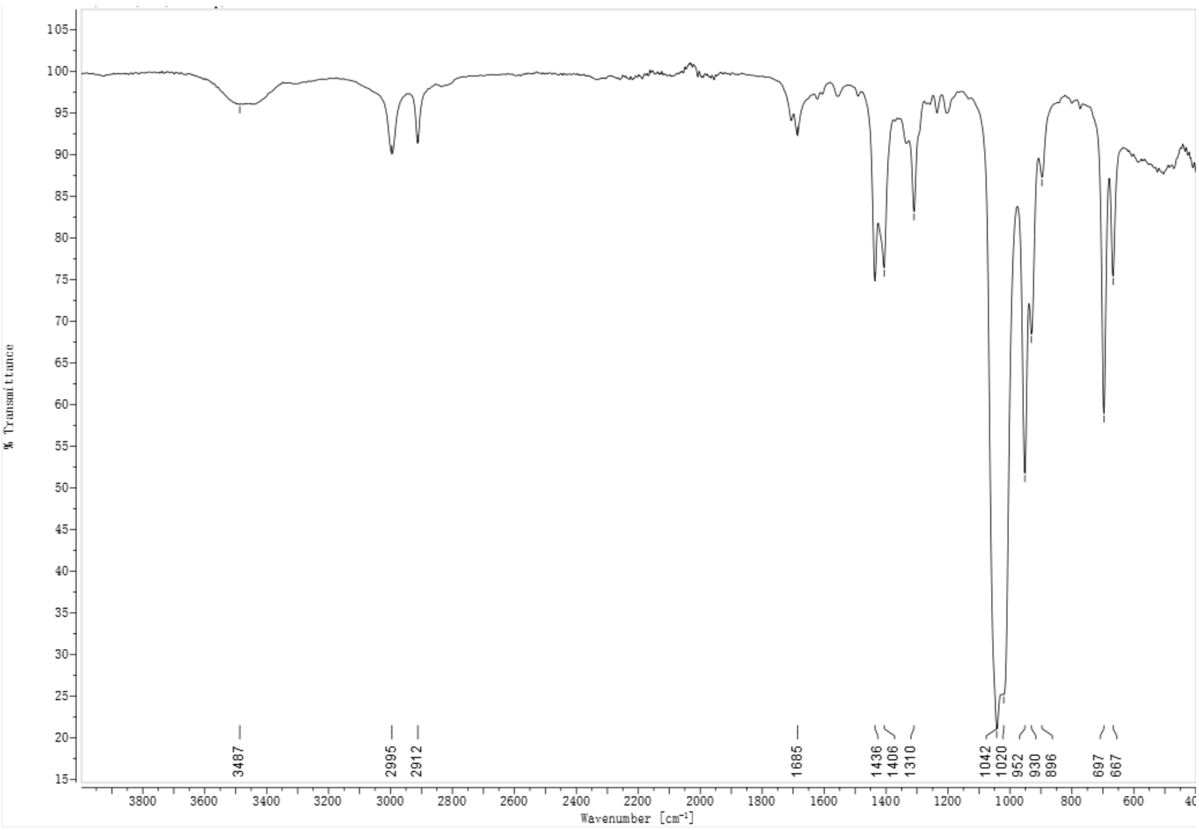
% Transmittance
Wavenumber [cm-1]
3487
2995
2912
1685
1436
1406
1310
1042
1020
952
930
896
697
667

HRMS of **SL1**:

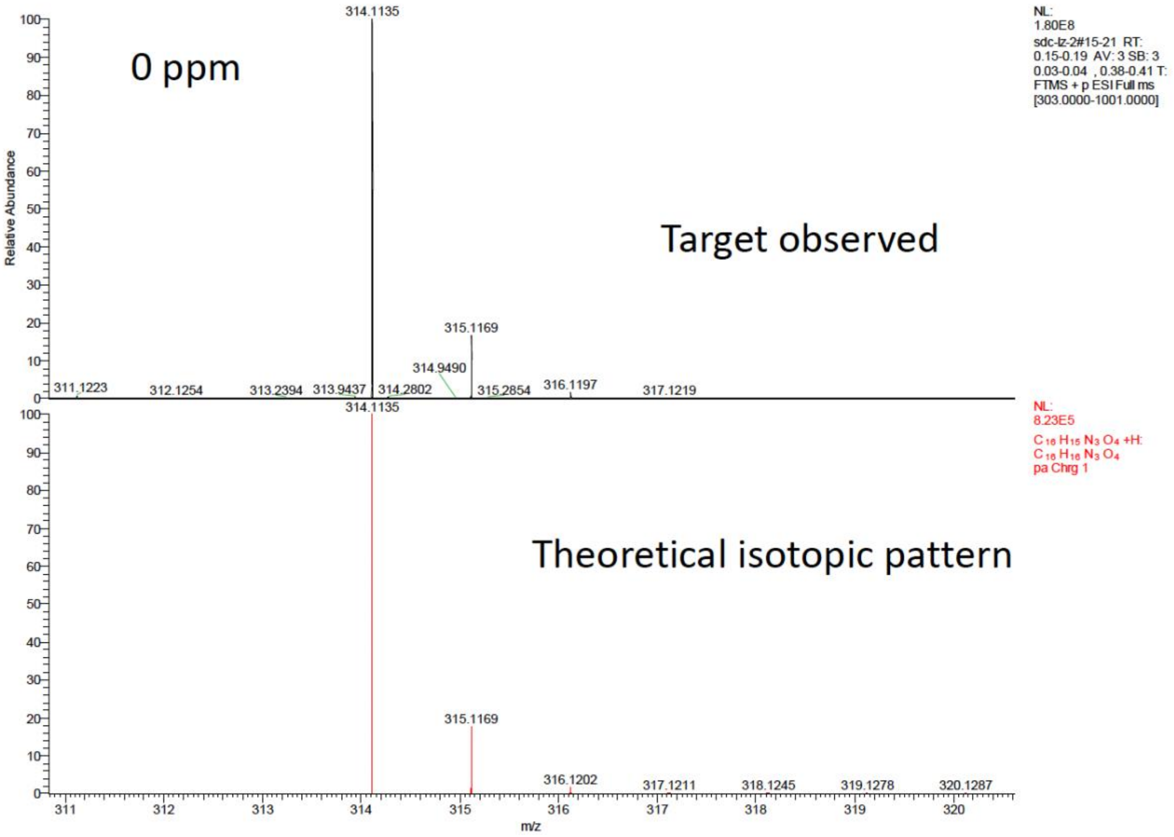
0 ppm
Target observed
Theoretical isotopic pattern
Relative Abundance
m/z
314.1135
315.1169
314.9490
311.1223
312.1254
313.2394
313.9437
314.2802
315.2854
316.1197
317.1219
316.1202
317.1211
318.1245
319.1278
320.1287
NL:
1.80E8
sdc-lz-2#15-21 RT:
0.15-0.19 AV: 3 SB: 3
0.03-0.04 , 0.38-0.41 T:
FTMS + p ESI Full ms
[303.0000-1001.0000]
NL:
8.23E5
C 16 H15 N3 O4 +H:
C 16 H16 N3 O4
pa Chrg 1
